# LGR5 targeting molecules as therapeutic agents for multiple cancer types

**DOI:** 10.1101/2022.09.01.506182

**Authors:** Hung-Chang Chen, Nico Mueller, Katherine Stott, Eilidh Rivers, Chrysa Kapeni, Carolin M Sauer, Flavio Beke, Stephen Walsh, Nicola Ashman, Louise O’Brien, Amir Rafati Fard, Arman Godsinia, Fadwa Joud, Olivier Giger, Inti Zlobec, Ioana Olan, Sarah J. Aitken, Matthew Hoare, Richard Mair, Eva Serrao, James D Brenton, Alicia Garcia-Gimenez, Simon E. Richardson, Brian Huntly, David R. Spring, Mikkel-Ole Skjødt, Karsten Skjødt, Marc de la Roche, Maike de la Roche

**Affiliations:** University of Cambridge, Cancer Research UK Cambridge Institute; Robinson Way, Cambridge CB2 0RE, UK; University of Cambridge, Department of Biochemistry; Tennis Court Road, Cambridge CB2 1QW, UK; University of Cambridge, Yusuf Hamied Department of Chemistry; Lensfield Road, Cambridge, CB2 1EW, UK; University of Cambridge, Department of Pathology; Tennis Court Road, Cambridge CB2 1QP, UK; Institute of Pathology, University of Bern; Murtenstrasse 31, CH-3008 Bern Switzerland; University of Cambridge, MRC Toxicology Unit; Tennis Court Road, Cambridge CB2 1QR, UK; Department of Histopathology, Cambridge University Hospitals, NHS Foundation Trust; Main Drive, Cambridge, CB2 0QQ, UK; University of Cambridge, Department of Haematology; Puddicombe Way, Cambridge CB2 0AW, UK; Rigshospitalet - University Hospital Copenhagen; Blegdamsvej 9, 2100 Copenhagen, Denmark; Institute of Immunology and Microbiology, University of Copenhagen; Blegdamsvej 3B, 2200 Copenhagen, Denmark; University of Southern Denmark; J.B. Winslows Vej, 5000 Odense C, Denmark

## Abstract

Leucine-rich repeat-containing G-protein receptor 5 (LGR5) has been characterised as a stem cell and cancer stem cell marker. Previous analyses of LGR5 transcript levels indicate high level expression discriminates malignancies such as colorectal cancer (CRC) and pre-B acute lymphoblastic leukaemia (pre-B ALL) from healthy tissues suggesting LGR5 protein expression may provide a molecular handle for prognosis and treatment.

We have developed highly specific, high affinity antibodies to the extracellular domain of human LGR5 (α-LGR5) that detect high LGR5 protein levels in colorectal cancer (CRC), hepatocellular carcinoma (HCC), and pre-B ALL. In contrast, there is low to undetectable levels of LGR5 protein in normal colon and rectal epithelia, liver, ovarian tissues, brain and immune cell types.

LGR5 is rapidly internalised from the plasma membrane and trafficked to intracellular vesicular compartments including lysosomes. Treatment of high LGR5-expressing CRC and pre-B ALL cancer cell lines with an antibody-drug conjugate version of α-LGR5 (α-LGR5-ADC) lead to effective cell killing at nanomolar concentrations. Interventional treatment of pre-B ALL tumours with α-LGR5-ADC *in vivo* led to rapid tumour attrition. We further demonstrated the therapeutic utility of humanised α-LGR5 by using the corresponding scFv fragment for the generation of α-LGR5 chimeric antigen receptors (CARs) and a Bispecific T cell Engager (BiTE). α-LGR5-CAR-NK cells were effective at killing LGR5-expressing cells while α-LGR5/α-CD3 BiTEs induce T cell activation and killing of NALM6 cells by cytotoxic CD8^+^ T cells.

Taken together, this study establishes α-LGR5-based therapeutic modalities that effectively discriminate and target CRC, HCC and pre-B ALL tumour cells.

**One Sentence Summary:** We generated novel antibodies against the cancer cell marker LGR5, validated diagnostic use in prioritizing specific cancer types for targeting, and developed antibody-based therapeutics.

## INTRODUCTION

Leucine-rich repeat-containing G-protein receptor 5 (LGR5) was identified as a target gene of oncogenic form of the Wnt signalling pathway in colorectal cancer cells (1). Subsequent work established Lgr5 expressing cells as functional stem cells in the intestinal epithelia (2), gastric epithelia (3), hair follicle (4), foetal mammary gland (5), nephrons in the developing kidney (6) and the regenerating liver (7).

LGR5 has attracted a great deal of therapeutic interest owing to its overexpression in malignancies such as CRC, hepatocellular carcinoma (HCC) (8), gastric and ovarian cancers (9), basal cell carcinoma (10), ER negative breast cancers (11), glioblastoma, (12) and certain B cell malignancies (13,14). In particular increased LGR5 transcript levels distinguish CRC tumours and intestinal adenocarcinomas from adjacent healthy tissue and specifically the invasive front of tumours and metastatic cells (reviewed in (15)). These LGR5 expression studies have measured transcript levels rather than protein levels.

Functionally, LGR5 expression in CRC cell lines is required for proliferation, migration, chemosensitivity, colony formation and *in vivo* transplantation ability (reviewed in (15)). *In vivo* studies using murine-engrafted human primary CRC cells and organoids have found that ablation of the LGR5 expressing cellular compartment, when treated with either a specific chemical toxin or chemotherapeutic agent, results in tumour regression (16,17). Interestingly, treatment withdrawal restores tumour proliferation, driven by cells that obligately acquire LGR5 expression. Overall, prognostic and functional studies of LGR5 expression in CRC and other malignancies support the intriguing notion that LGR5 expressing tumour cells are a promising therapeutic target for cancer treatment.

Antibody-based immune therapies harness the specificity, efficacy and versatility of antibodies for recognition of cell surface proteins overexpressed on cancer cells leading to targeted ablation. Antibodies have been deployed therapeutically for antibody-dependent cellular cytotoxicity (ADCC) against cancer cells, as antibody-drug conjugates (ADCs) for directing an attached toxin to cancer cells or as immune cell re-directing strategies that include chimeric antigen receptors (CARs) or the use of bispecific immune cell engagers (e.g. BiTEs) that connect cytotoxic immune cells to cancer cell targets (18–20). Key advances in the immune therapy field will require the identification of new molecular targets on tumour cells and the development of high-quality antibodies to direct immune system components or cytotoxic agents.

While there is clinical scope for use of LGR5 antibodies in immune therapies, there are few, if any, commercial antibodies that have undergone robust validation, hampering therapeutic development. Here we report the development of highly specific monoclonal α-LGR5 antibodies that are compatible with a range of detection and therapeutic applications. Using α-LGR5 as a discovery tool, we have determined that LGR5 protein rapidly internalises from the cell surface and is trafficked to intracellular vesicles that overlap with the lysosomal compartment. Our census of LGR5 protein expression on healthy human tissues and malignancies establish low to undetectable level of LGR5 in healthy tissues and specific overexpression of the protein in CRC, hepatocellular carcinoma and pre-B ALL tumours. An antibody drug conjugate (ADC) version of the α-LGR5 antibody eradicates human cell line models of LGR5-expressing CRC and pre-B ALL *in vitro* and demonstrates high efficacy in targeting a pre-B ALL tumour model *in vivo*. Finally, CAR and BiTE versions of α-LGR5 effectively direct cytotoxic immune cell killing of LGR5-expressing cancer cells.

Taken together, LGR5 is a promising tumour-specific biomarker in a subset of cancers that can be detected and targeted by our newly developed LGR5 antibody-based modalities.

## RESULTS

### Generation and validation of antibodies against LGR5

For the monoclonal antibody production, we immunized mice with the N-terminal 100 amino acids of the human LGR5 extracellular domain (**Suppl. Fig. 1A**). Coupling of the antigen to diphtheria toxiod was necessary to initiate an effective immune response. B cell fusions yielded 18 hybridoma clones which were tested against transgenic versions of human and murine LGR family members expressed in HEK293T cells. A common sequence coding for an N-terminal influenza hemagglutinin (HA) epitope tag and a C-terminal extension consisting of the fusion to the vasopressin V2 receptor C-terminal tail for enhanced protein stability fused to eGFP was added to all human and murine LGR family transgenes (**Suppl. Fig. 1B**) (21). We also created a version of human LGR5 with Gly1Ser and Val8Ala substitutions to match the corresponding *cynomolgus* LGR5 N-terminus (cLGR5); the N-terminal 100 amino acid residues of cLGR5 are otherwise identical to the human LGR5 protein (hLGR5). Western blot analysis of lysates derived from HEK293T cells expressing the various LGR transgenes demonstrated immunoreactivity of the four hybridoma clones (clones 1-4) exclusively to human and *cynomolgus* LGR5 (**Fig. 1A, Suppl. Fig. 1C**). We did not observe any immunoreactivity of the hybridoma clones to non-transfected HEK293T cells, as these cells do not express LGR5 in the absence of pathway activity.

**Fig. 1.**
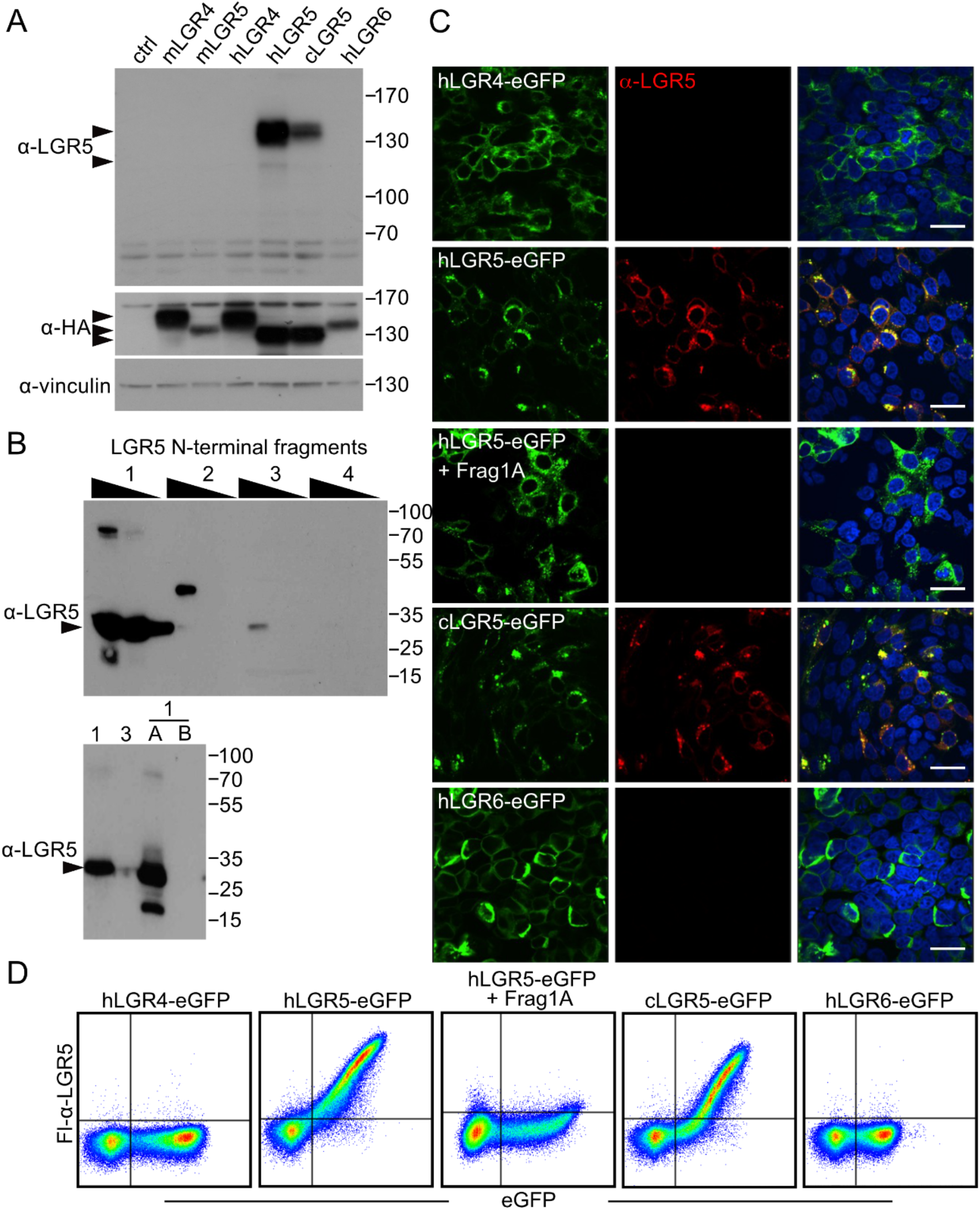
Validation of novel α-LGR5 antibody clones. A. Representative western blot analysis of HEK293T lysates expressing LGR family transgenes probed with α-LGR5 clone 2 and antibodies raised against HA and vinculin. Arrowheads on the side of the top panel indicate expected position of hLGR5-eGFP (upper) and endogenous LGR5 (lower). Arrowheads on the side of the middle panel indicate the expected sizes of the HA-tagged LGR5 family transgenes. B. Epitope mapping of α-LGR5 using fragments of the 100 amino acid antigen sequence (**Suppl. Fig. 1A**) traces the epitope of hybridoma clone 2 to Frag1A, the N-terminal 15 amino acids of the mature human LGR5. C. FL-α-LGR5 detection of LGR5 (red) in HEK293T cells overexpressing eGFP-fused human LGR4-6 transgenes (hLGR4-6) and *cynomolgus* LGR5-eGFP (cLGR5-eGFP; green). In the third row of panels, pre-incubation of FL-α-LGR5 with Frag1A, attenuates the signal for detection of hLGR5-eGFP. Blue, DAPI fluorescence showing nuclei. Scale bars, 10 μM. D. Flow cytometric analysis of HEK293T cells expressing hLGR family transgenes using FL-α-LGR5 and eGFP for detection. *Second panel* - pre-incubation of FL-α-LGR5 with Frag1A abrogates the signal detecting hLGR5-eGFP expressing cells.

### All α-LGR5 antibody clones bind a common epitope at the LGR5 N-terminus

The amino acid sequences of the complementary determining regions (CDRs) of the light and heavy chains of the four immune reactive α-LGR5 antibodies are highly conserved, displaying variation at only four positions (**Suppl. Fig. 1D**). To determine whether the four α-LGR5 clones bound a common epitope, we generated four overlapping fragments, approximately 35 amino acids in length (Fragments 1-4; **Suppl. Fig. 1A**), using the RAD-display fusion system (22). Western blot analysis indicated that all four α-LGR5 clones were specifically immunoreactive to Fragment 1 (**Fig. 1B; Suppl. Fig. 1E**). We further refined the epitope sequence to Fragment 1A (Frag1A), a 15 amino acid sequence at the N-terminus of the LGR5 protein (**Suppl. Fig. 1A**) that was bound by all four α-LGR5 clones (**Fig. 1B, Suppl. Fig. 1E**). None of the clones bound to the adjacent, overlapping, 15 amino acid Frag1B (**Fig. 1B, Suppl. Fig. 1A**). Notably, the sequence of the Frag1A epitope diverges substantially in sequence from the corresponding region in human LGR4/6 and mouse LGR4/5 and by a single conserved residue in the cLGR5 sequence (**Suppl. Fig. 1F**) explaining why clones 1-4 are specific to the human and *cynomolgus* LGR5.

Binding affinities between the α-LGR5 antibody clones and Frag1A and Frag1B were determined by bio-layer interferometry measurements using the Octet platform. The bidentate nature of the two arms of the antibody clones resulted in slow dissociation due to re-binding. Therefore, accurate determination of the *Kd* values required inclusion of 10 μM of a competitor peptide corresponding to the Frag1A sequence during the binding assay. We observed high affinity binding between Frag1A and the antibody clones with *Kd* values between 0.76 and 1.4 nM (**Table 1**). No detectable binding was observed between the antibody clones and Frag1B.

**Table 1.**
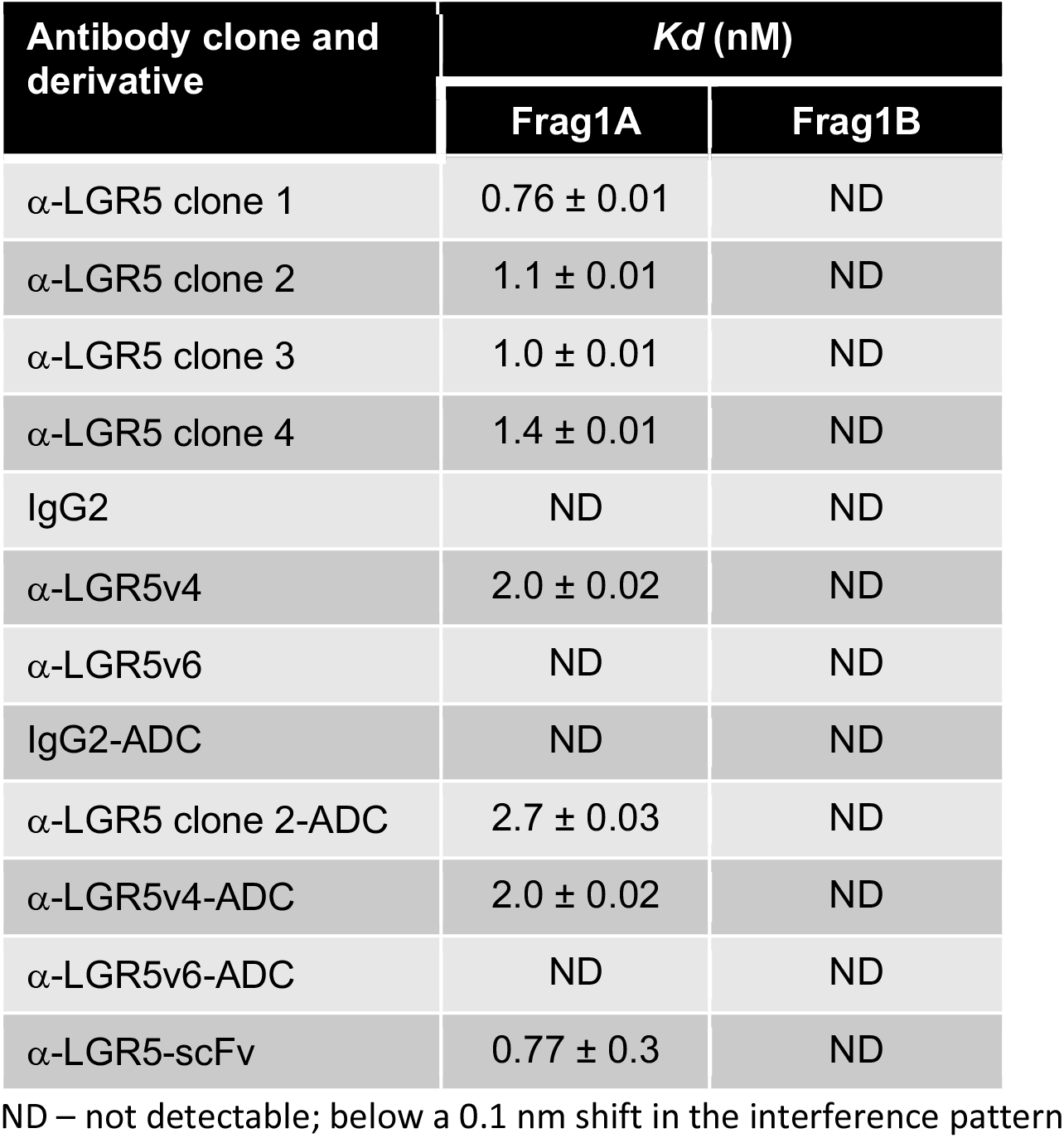
Binding affinities for murine, humanized and ADC-modified α-LGR5 antibodies.

Because of the proximity of the LGR5 Frag1A sequence to amino acids that mediate binding to R-spondin family ligands (23), we determined whether antibody binding interfered with Wnt pathway activity in LGR5 expressing cells, measured using the TopFlash assay (24). We treated HEK293T cells expressing either hLGR5-eGFP or eGFP controls with Wnt and R-spondin1, in the presence of greater than 10-fold molar excess of either antibody hybridoma clone 2 (α-LGR5; relative to R-spondin1) or murine IgG1 as control. There was no significant difference in Wnt/R-spondin1-activated pathway activity between IgG1 or α-LGR5 treated cells (**Suppl. Fig. 1G)** indicating antibody binding to LGR5 does not interfere with R-spondin1 binding. Taken together, the α-LGR5 antibody binds with high affinity to the N-terminus of human and cLGR5 at a site that does not interfere with binding of R-spondin ligands.

### α-LGR5 antibodies bind human and cynomolgus LGR5 but no other LGR family members

To determine antibody specificity in detecting cellular LGR5 expression, we overexpressed the LGR transgenes in HEK293T cells, fixed the cells in formaldehyde, and probed expression with murine α-LGR5 coupled to Dylight650 fluorophore (Fl-α-LGR5) by immunofluorescence. We did not observe fluorescence signal from Fl-α-LGR5 in the absence of transgenic expression; however, robust co-localisation of Fl-α-LGR5 and over-expressed hLGR5-eGFP was detected and the Fl-α-LGR5 signal was abrogated by pre-incubation of the antibody with Frag1A (**Fig. 1C**). We could also detect the over-expressed cLGR5 transgene with Fl-α-LGR5 in HEK293T cells but not the over-expressed human LGR4, LGR6 or murine LGR4 and LGR5 transgenes (**Fig. 1C; Suppl. Fig. 1H**). Flow cytometric analysis of live HEK293T cells overexpressing human, murine, and *cynomolgus* version of LGR family members yielded identical results - hLGR5-eGFP and cLGR5-eGFP overexpressing HEK293T cells were detected by Fl-α-LGR5, but not cells overexpressing the other LGR family transgenes (**Fig. 1D**; **Suppl. Fig. 1I**). Importantly, the signal for cell surface hLGR5-eGFP-expressing cells was abrogated by pre-incubation of Fl-α-LGR5 with Frag1A but not Frag1B (**Suppl. Fig. 1I**).

Taken together, our α-LGR5 antibody specifically detects overexpressed hLGR5 and cLGR5 by western blot and immunofluorescence as well as flow cytometry.

### Census of LGR5 expression levels in healthy tissues and cancers

Previous studies have established high LGR5 transcript levels in CRC and some other cancers (25,26) raising the possibility of using α-LGR5 as a cancer biomarker or developing immune-based strategies for therapeutic targeting. We carried out our own comprehensive census of LGR5 transcript levels across 33 cancer types using datasets extracted from the TCGA database (**Suppl. Fig. 2A**). We assigned 11 of the cancer types as ‘high LGR5 expressors’, defined as cancers with more than 70% of component cases harbouring LGR5 expression levels greater than the pan-cancer median; these included head and neck, pancreatic, oesophageal, adrenal gland, brain, stomach, ovarian, uterine, colon, endometrial, and rectal cancers (**Suppl. Fig. 2A**). For selected high LGR5 expressors, we compared LGR5 expression to matched healthy tissue. Brain cancer, ovarian cancer and uterine carcinosarcoma were excluded from this analysis owing to insufficient data on healthy tissues in the TCGA database. Significant increases in LGR5 transcript expression were apparent in malignancies of the head and neck, uterine endometrial, stomach, colon and rectum (**Suppl. Fig. 2B**). The exceptions were liver cancer (hepatocellular carcinoma; HCC), where LGR5 expression was depressed relative to healthy tissue, and adrenal gland cancer, in which no significant difference in LGR5 expression was observed in comparison to healthy tissues.

**Fig. 2.**
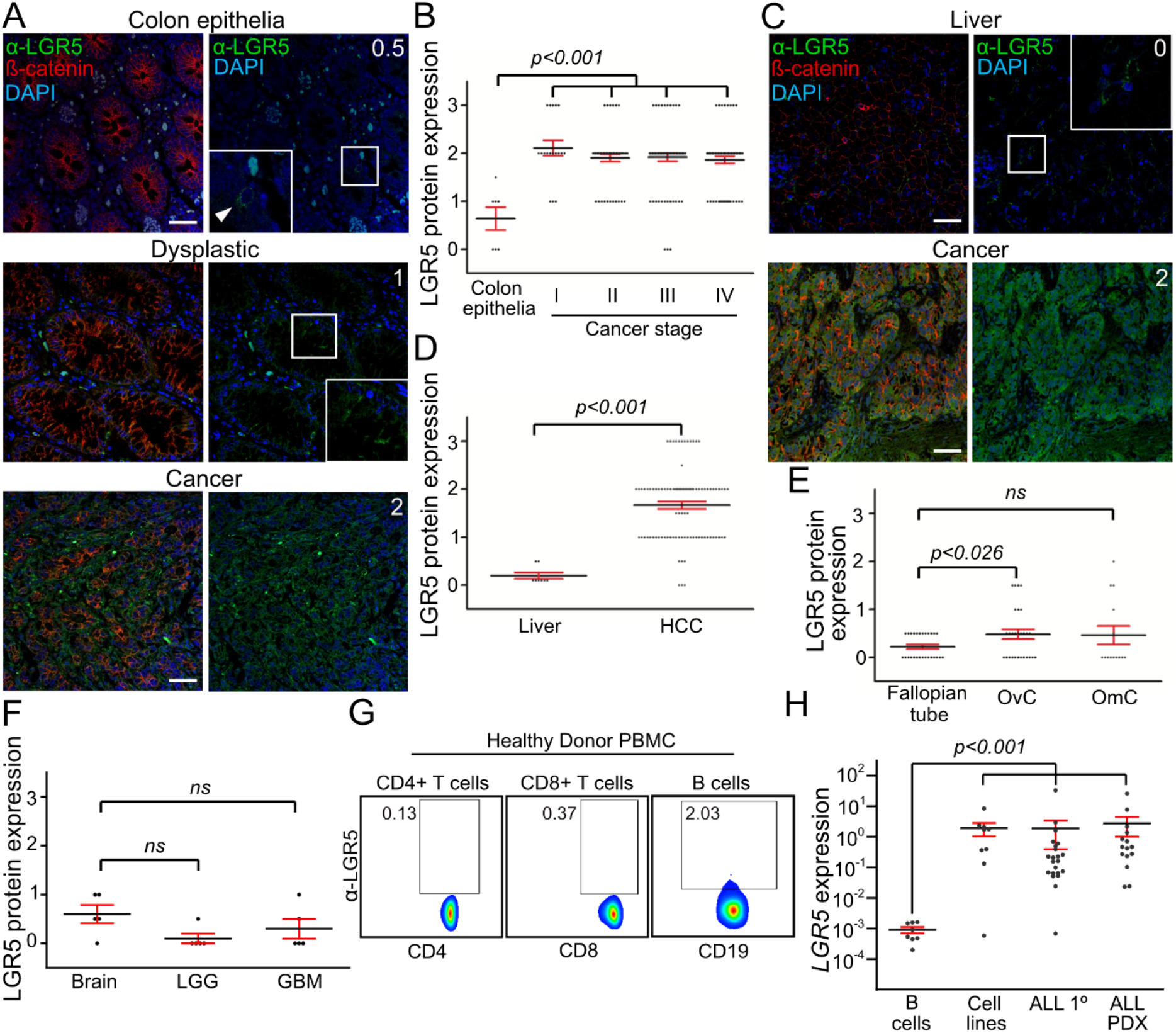
Census of healthy tissues and cancers for LGR5 expression levels. A. Sections from a CRC tumour resection showing LGR5 and β-catenin expression in - *top panels*, normal tissue; *middle panels*, dysplastic tissue; and *lower panels*, CRC. Blue, DAPI fluorescence showing nuclei. White number in the top right corner of the lower set of panels (showing LGR5 expression and nuclei staining) corresponds to relative levels of LGR5 protein using the scoring criteria applied to all TMAs (see text). Scale bar, 40 μM. B. Relative LGR5 protein expression quantified in healthy colon epithelia and CRC tumour stages I-IV scored. Each dot represents a single scored biopsy on the Bern TMA. Level of significance, *p-value*, determined by two-tailed t-tests comparing LGR5 expression for colon epithelia with each of the four tumour stages. C. Immunofluorescence using Fl-α-LGR5 and an antibody to β-catenin for - *top panels*, a healthy liver sample; and bottom *right panels*, a sample from the Cambridge HCC TMA. Numbers in white correspond to scored LGR5 expression levels using the criteria for evaluating the TMA. Arrowhead, cell clusters within the tumour with high levels of cortical β-catenin and low levels of LGR5 levels. White numbers represent scored values for relative LGR5 protein expression. Scale bar, 40 μM. D. Quantitation of LGR5 protein expression levels in 8 healthy liver resections (Liver) and biopies from the Cambridge HCC TMA (HCC). Level of significant difference, *p-value*, between the sample sets was determined by two-tailed t-test. E. Quantitation of LGR5 expression levels in biopsies of healthy Fallopian tube, ovarian cancer cases (OvC) and omentum cancer cases (OmC) comprising the Cambridge ovarian cancer TMA. Levels of significance in LGR5 expression between Fallopian tube and OvC samples, *p-value*, was calculated using two-tailed t-test. There were no significant increases (*ns*) in LGR5 expression between Fallopian tube and OmC sample sets. F. Quantitation of LGR5 expression levels in samples from the Cambridge Brain cancer TMA - healthy brain tissue (Brain), low grade glioma (LGG) and glioblastoma (GBB). There were no significant increases (*ns*) in LGR5 expression between Brain, LGG or GBB samples sets, determined by two-tailed t-test. G. Representative example of relative LGR5 protein expression levels in CD4^+^ T cells, CD8^+^ T cells and CD19^+^ B cells from healthy donor PBMCs determined by flow cytometry. A small number of B cells, <3% of the total population, express low levels of LGR5. H. *LGR5* transcript levels normalised to *TBP*, measured by quantitative RT-PCR, in healthy donor B cells (B-cells; 10 samples), B-ALL cell lines (Cell lines; 9 lines), CD19-enriched cell populations from primary B-ALL cases (ALL primary; 15 samples) and CD19-enriched populations from B-ALL tumour cells maintained as PDX models (ALL-PDX 22 samples). Level of significance, *p-value*, was determined by two-tailed t-tests comparing LGR5 expression in healthy donor B cells and the other sample sets at the *, *p<0*.*001* level of significance. LGR5 transcript levels for the individual samples are shown in **Suppl. Fig. 2L**.

Experimental differences in the generation of composite TCGA datasets may confound direct inter-tissue/tumour comparisons of LGR5 expression levels. Moreover, direct correlation between mRNA levels and functional protein levels is often inaccurate. We therefore used α-LGR5 to determine LGR5 protein levels in several cancer types by indirect immunofluorescence and a compatible antibody to β-catenin (α-β-catenin) to distinguish epithelial cells.

For CRC, we directly compared LGR5 expression levels in three individual biopsies each containing regions of healthy colon epithelia, dysplastic epithelia and cancer. In the healthy colon epithelia, LGR5 expression was confined to a small number, <1%, of β-catenin positive cells and consisted of intracellular puncta with no indication of plasma membrane staining (**Fig. 2A, Suppl. Fig. 2C**). In all three cases, both the number of LGR5-expressing cells and the cellular levels of LGR5 protein in puncta increased with disease progression from dysplastic epithelia to cancer.

To validate our findings, we probed the highly annotated Bern CRC tumour microarray (TMA) with α-LGR5 and α-β-catenin antibodies (27). Healthy colon epithelia and CRC tumours were scored for levels of LGR5 expression on a scale of 0-3 in β-catenin positive cells: a score of 0 corresponds to the absence of any signal; 1 corresponds to low signal in less than 10% of cells; 2 is either higher signal for LGR5 and/or signal in greater than 20% of cells; and 3 is higher signal for LGR5 in greater than 20% of cells. Of the 213 scored biopsies on the TMA, LGR5 overexpression was apparent in 80% of the tumour samples; however, we did not observe significant differences between tumour stages (**Fig. 2B**) and other phenotypic metrics evaluated for the Bern CRC TMA that include patient age and gender groups, tumour location or the presence of microsatellite instability.

Our LGR5 transcriptional analysis revealed no elevation in hepatocellular carcinoma (HCC) contrasting previous studies of the malignancy (25,26). We therefore probed LGR5 protein levels using a TMA consisting of cores from the centre and edge of 103 HCCs, along with control healthy liver biopsies taken from five women having undergone resections for inflammatory adenoma (3 cases) or focal nodular hyperplasia (2 cases) (**Fig. 2C, D; Suppl. Fig. 2D**). The control liver biopsies displayed undetectable or very low levels of LGR5 protein localized to the hepatocyte cell cortex, overlapping with β-catenin (**Fig. 2C; Suppl. Fig. 2D**). Of note, we also observed high levels of cortical β-catenin, approximately 5-fold higher, localized to cell clusters representing bile ducts (**Suppl. Fig. 2D**). Importantly, we observed significantly elevated levels of LGR5 in the HCC cases on an HCC TMA for 65% of the cases (**Fig. 2D**; **Suppl. Fig. 2D**). β-catenin was overexpressed in 82% and the concordance in overexpression between the two proteins was 64% (**Suppl. Fig. 2E**). As with the control liver, we observed cell clusters within the HCC tumours with high levels of cortical β-catenin where LGR5 levels were consistently 3-5-fold lower than in adjacent cells (**Suppl. Fig. 2D**). LGR5 expression was also able to identify tumoral sub-groups: the non-proliferation class of HCC is dominated by activating mutations in the Wnt pathway component β-catenin (or in rare cases AXIN1), lack of AFP expression and relatively low proliferation rates (28). Across the cohort, we found negative correlations between LGR5 expression and serum AFP at time of transplant (**Suppl. Fig. 2F**) and tumour cell proliferative fraction of by Ki67 staining (**Suppl. Fig. 2G**). Therefore, LGR5 expression was elevated in tumours with clinical and molecular features of the non-proliferative HCC sub-class.

Pancreatic expression of LGR5 transcript was the lowest of the healthy tissues analysed and showed increased levels in pancreatic cancer. However, of the 5 matched healthy and tumour samples we analysed, we failed to see a consistently increased LGR5 protein expression in malignant tissues (**Suppl. Fig. 2H**).

Our transcription data ranked ovarian cancer as one of the highest LGR5 expressing cancers. However, owing to lack of data for fallopian tubes in the TCGA database, the presumptive tissue of origin for ovarian cancer, we were unable to determine whether this represents malignancy specific LGR5 overexpression. We therefore probed 24 fallopian tube biopsies with α-LGR5 and α-β-catenin alongside a TMA containing 28 ovarian cancer and 14 omentum cancer cases (**Suppl. Fig. 2I**). We did not observe a significant increase in LGR5 protein levels in ovarian and omentum cancer cases (**Fig. 2E**). β-catenin was expressed at approximately equal levels at the cortex of all epithelial cells and cancer cells (**Suppl. Fig. 2J**). We did not detect appreciable LGR5 protein levels in the β-catenin positive epithelia cells except in very rare instances where we observed individual cells expressing LGR5 localized to intracellular puncta (less than <0.1% cells; **Suppl. Fig. 2I**).

Finally, we used α-LGR5 to probe brain cancers (glioblastoma and low-grade glioma; LGG) using a TMA containing malignant and healthy brain biopsies. β-catenin levels were generally low and we did not detect LGR5 expression in healthy brain tissue and no significant upregulation of LGR5 protein was observed in glioblastoma and LGG (**Fig. 2F; Suppl. Fig. 2J**).

A previous study has established a functional role for LGR5 in murine B cell development and correlated LGR5 overexpression with a poor clinical outcome in human pre-B ALL (13,14). Thus, we extended our LGR5 protein expression analyses to immune cells and leukaemias. Flow cytometric analysis of human peripheral blood mononuclear cells (PBMCs) using α-LGR5 showed no expression of LGR5 in CD4^+^ and CD8^+^ T cells; however, we identified a small proportion (approximately 2%) of CD19^+^ B cells with potentially low levels of LGR5 expression **(Fig. 2G**). To determine whether LGR5 levels are increased in leukaemias, we quantified LGR5 expression in patient-derived, primary acute lymphoblastic leukaemia (ALL), ALL patient-derived xenograft (PDX) models and pre-B ALL cell lines. In the patient derived samples (ALL and PDX), we found that 29 out of the 39 cases (74%) showed a significant increase in LGR5 mRNA levels with two cases displaying greater than 100-fold increases compared with healthy B cell controls (**Fig. 2H; Suppl. Fig. 2K**). We also observed significantly elevated LGR5 transcript levels in 8 out of 9 pre-B ALL cell lines.

Taken together, our analyses of LGR5 transcript and protein levels in healthy tissues and malignancies establish that: (i) LGR5 overexpression is endemic to a discrete set of cancer types; (ii) there is a substantial window of LGR5 expression between CRC, HCC and ALL and healthy tissue; and (iii) the use of specific and well validated antibodies such as α-LGR5 is essential for establishing functional protein levels in cancer.

### α-LGR5 detects endogenous LGR5 expression

We next determined cellular expression levels of LGR5 protein in three of the pre-B ALL cell lines; NALM6, REH and 697. LGR5 can be detected by western blot in pre-B ALL cell lines (**Fig. 3A**) where protein levels in the three cell lines accurately reflect relative transcription levels (**Suppl. Fig. 2J**) - NALM6 cells harbour higher protein levels than REH cells and 697 cells express very low levels of LGR5.

**Fig. 3.**
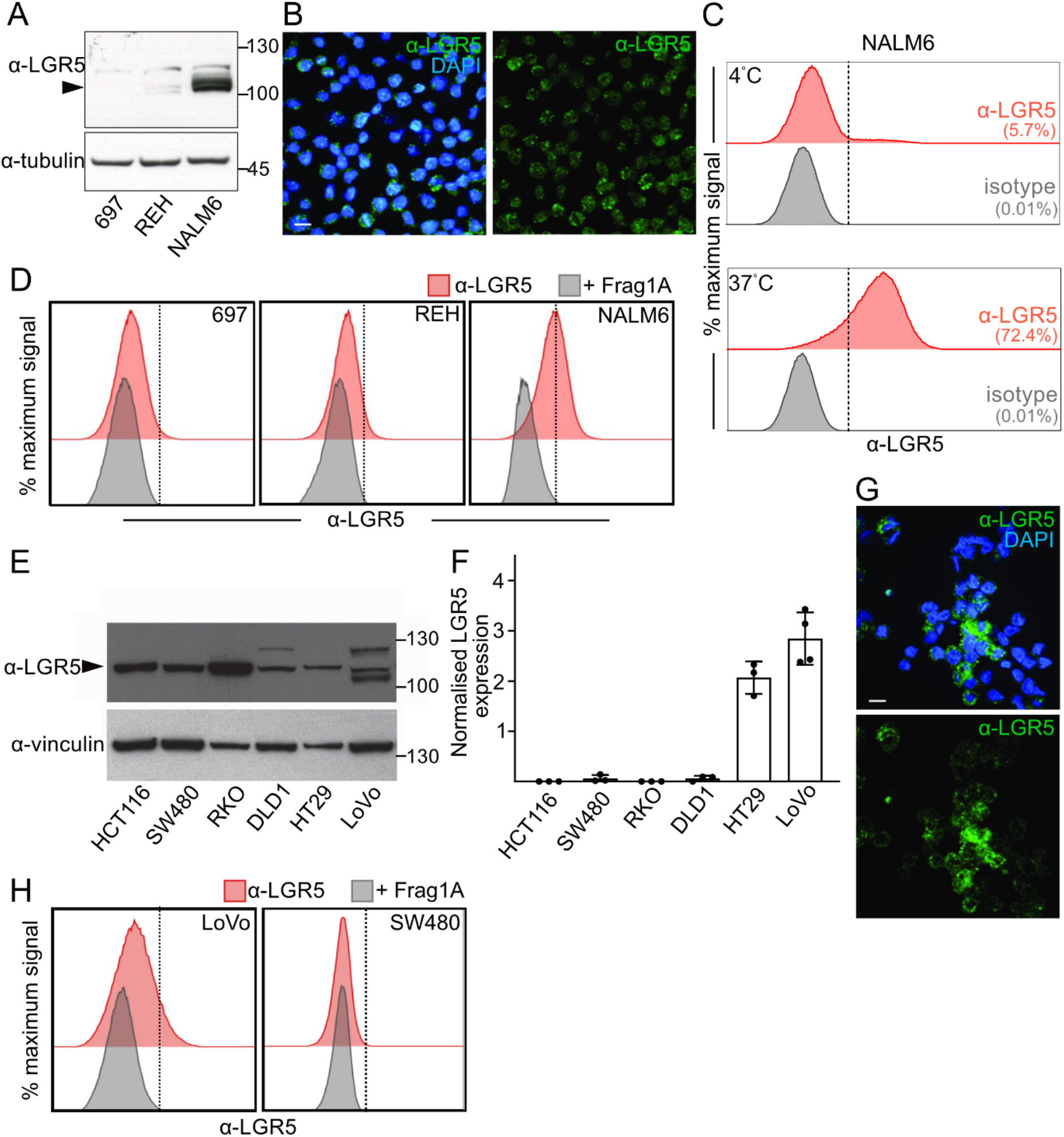
Characterisation of LGR5 expression in NALM6 and LoVo cell lines. A. Western blot analysis of LGR5 protein levels in lysates from pre-B ALL cell lines. B. Indirect immunofluorescence of NALM6 cells using α-LGR5. Scale bar, 5 μM. C. Flow cytometric detection of Fl-α-LGR5 (red histograms) or fluorescent isotype control (grey histograms) association with NALM6 cells after 1 hour incubation at 4°C (top panel) or 37°C (bottom) panel. Numbers represent percentage of cells with detectable fluorescence. D. Flow cytometric detection of Fl-α-LGR5 (red histograms) association with 697, REH or NALM6 cells after 1 hour incubation at 37°C. Grey histograms – analysis with Fl-α-LGR5 pre-incubated with Frag1A. E. Representative western blot analysis of LGR5 protein in CRC cell lines using α-LGR5 and an antibody to vinculin as loading control. F. Relative *LGR5* transcript levels normalised to *TBP* in CRC cell lines measured by qRT-PCR. Error bars indicate standard deviation (SD) for 3 biological replicates except for LoVo cells, 4 biological replicates. G. Indirect immunofluorescence of LGR5 in LoVo cells using α-LGR5. Scale bar, 5 μM. H. Flow cytometric analysis of LoVo and SW480 cells that have been incubated with Fl-α-LGR5 (red plots) or with Fl-α-LGR5 pre-incubated with Frag1A (grey plots) at 37°C for 1 hour.

The intracellular distribution of LGR5 was determined by immunofluorescence using α-LGR5. LGR5 localises to intracellular puncta with no observable association with the cell periphery (**Fig. 3B**). Flow cytometry analysis of NALM6 cells was carried out with a version of α-LGR5 labelled with the Dylight TM650 fluorophore (Fl-α-LGR5). We conducted these experiments using 1-hour incubations of NALM6 cells with Fl-α-LGR5 at 4°C and detected very low levels of LGR5 on the cell surface (**Fig. 3C**). However, when Fl-α-LGR5 and NALM6 cell were incubated for 1-hour at 37°C, the entire NALM6 population stained positive for LGR5 (**Fig. 3C**) suggesting antibody internalisation. We did not detect association/internalisation of the fluorescent IgG1 control. Control flow cytometry experiments were carried out using the low LGR5-expressing REH and 697 cell lines by incubation with Fl-α-LGR5 at 37°C for 1 hour. Importantly, we were only able to detect slight Fl-α-LGR5 internalisation by 697 and REH cells relative to by NALM6 cells (**Fig. 3D**), consistent with our previous quantification of cellular LGR5 protein and transcript levels. We did not observe any staining by Fl-α-LGR5 when the antibody was pre-incubated with Frag1A. Our data indicates that LGR5 is predominantly expressed in intracellular puncta and is internalised from a small plasma membrane protein pool.

The LoVo colon cancer cell line has previously been shown to express sufficient levels of LGR5 for antibody detection (25,26) and indeed, we found that amongst a panel of five additional colorectal cancer cell lines (HCT116, SW480, RKO, DLD1 and HT29), LoVo cells expressed high levels of LGR5 protein (**Fig. 3E**) and the highest LGR5 transcript levels (**Fig. 3F**). Immunofluorescent detection of LGR5 in LoVo cells using Fl-α-LGR5 was consistent with NALM6 cells; LGR5 expression is confined to intracellular puncta and not observable on the cell surface (**Fig. 3G**). Similar to NALM6 cells, flow cytometric detection of live LoVo cells required pre-incubation of Fl-α-LGR5 for 1 hour at 37°C indicating LGR5 internalisation (**Fig. 3H**). In contrast, we do not observe LGR5 internalisation by SW480 cells, perhaps the consequence of lower protein expression levels (**Fig. 3H**). We note that the punctate intracellular distribution of LGR5 we observe in LoVo and NALM6 cells matches that in CRC and HCC tumour cells as well as specific LGR5 expressing cells in colon and rectal epithelia, liver and fallopian tubes (**Fig. 2A and C, Suppl. Fig. 2C and D**). We hypothesise that the steady-state intracellular distribution of LGR5 reflects a rapid internalisation from the plasma membrane and trafficking to intracellular puncta.

### Rapid internalization of α-LGR5 antibodies by LGR5-overexpressing cell lines

To determine the kinetics of LGR5 internalisation we treated LGR5-eGFP overexpressing HEK293T cells with Fl-α-LGR5 - within 5 minutes Fl-α-LGR5 was internalised and localised to puncta juxtaposed to the cell surface. Ultimately the fluorescent signal from Fl-α-LGR5 associated entirely with the intracellular LGR5-eGFP puncta over the course of 45 minutes (**Fig. 4A**). By contrast, HEK293T cells that were transfected with LGR4-eGFP failed to bind or internalize FL-α-LGR5 (**Fig. 4A**).

**Fig. 4.**
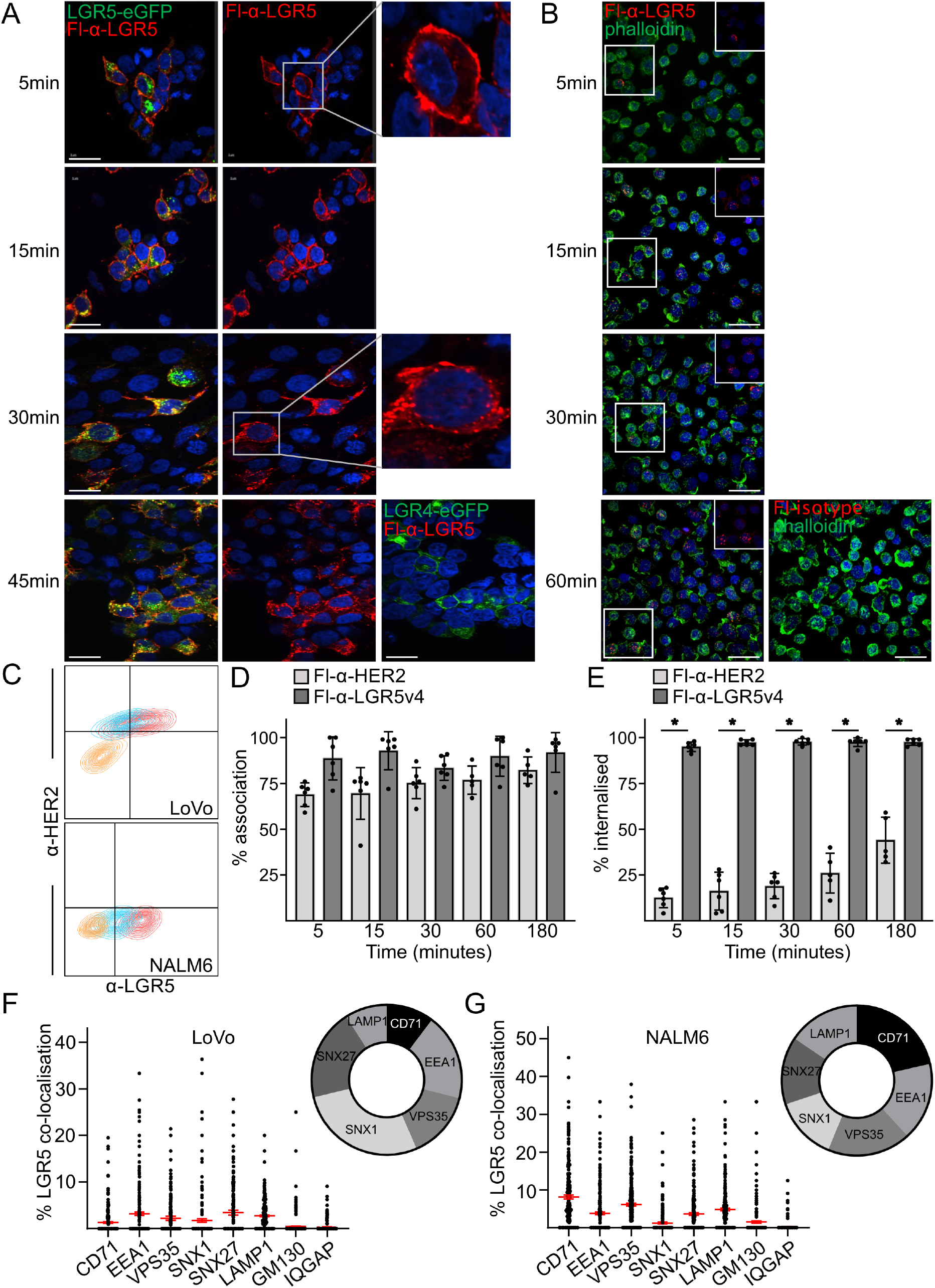
Rapid internalisation of cell surface LGR5. A. Time course of FL-α-LGR5 internalisation by LGR5-eGFP expressing HEK293T cells. *Top right panel* – enlarged image showing association of FL-α-LGR5 with the cell periphery after 5 minutes, followed by co-localisation with the internal LGR5-eGFP associated puncta within 30 minutes (*middle right panel*). *Bottom right panel* – no association or internalisation of FL-α-LGR5 for cells expressing LGR4-eGFP. Scale bar, 10 μM. B. Time course of Fl-α-LGR5v4 (red) internalisation by NALM6 cells detected by immunofluorescence. At the timepoints, fixed cells were probed with fluorescent (Alexa488) phalloidin (green) and Hoechst (blue). Bottom panels, Fl-α-LGR5v4, substituted with Fl-α-LGR5v6 (red) as isotype control. In the bottom right panel, the signal from the fluorescent Alexa488 phalloidin has been omitted. Scale bar, 10 μM. C. Flow cytometric analysis of LoVo and NALM6 cells after 60 minutes incubation with Fl-IgG1 (orange), and Fl-α-HER2 (blue) with either Fl-α-LGR5v4 (red). Each panel is a composite of the individual analyses in **Suppl. Fig. 4B**. D. Time course of Fl-α-LGR5v4 and Fl-α-HER2 association with LoVo cells. Percent association was scored as fraction of cells associated with fluorescent signals from Fl-α-HER2 (light grey bars) or Fl-α-LGR5v4 (dark grey bars) at the indicated timepoints per field of view (example images shown in **Suppl. Fig. 4D**). Data points are average of 6 individual scoring experiments, each measuring at 80 to 200 cells over for 2 independent experiments. Error bars indicate SD amongst scoring experiments. There were no significant differences between Fl-α-LGR5v4 or Fl-α-HER2 association with LoVo cells or amongst any of the timepoints. E. Time course of percent internalisation of Fl-α-LGR5v4 and Fl-α-HER2 by LoVo cells. Internalisation data scores 6 experimental counts of 80-200 cells over 2 independent experiments. Error bars indicate SD. Significant differences in internalisation between Fl-α-LGR5v4 or Fl-α-HER2 are at the *, *p<0*.*001* level of significance, determined by two-tailed t-test, for the 5- to 180-minute timepoints. F. Co-localisation between internalized FL-α-LGR5v4 puncta and markers of various intracellular compartments and IQGAP1 in LoVo cells. Co-localisation data were derived from automatic scoring of puncta in images of internalised FL-α-LGR5v4 and intracellular markers detected by indirect immunofluorescence and is the composite of a minimum of 200 cells over 2 individual experiments. *Right* - wheel graph indicating fractional association of internalised FL-α-LGR5v4 with specific intracellular vesicle markers in LoVo cells. Representative images are shown in **Suppl. Fig. 4F**. Error bars SD, data is the product of two independent experiments. G. Co-localisation between internalized FL-α-LGR5 puncta and markers of various intracellular compartments and IQGAP1 in NALM6 cells, scoring as above. *Right* - wheel graph showing fractional association of internalised FL-α-LGR5v4 with specific intracellular vesicle markers in NALM6 cells. Representative images used for co-localisation datasets are shown in **Suppl. Fig. 4G**. Data is derived from two independent experiments.

The rate of LGR5 internalisation was monitored for NALM6 and LoVo cells and both displayed identical kinetics to internalisation by LGR5-eGFP-overexpressing HEK293T cells: Fl-α-LGR5-associated intracellular puncta were evident within 5 minutes increasing to a maximum number of puncta by 30 minutes for both NALM6 (**Fig. 4B**) and LoVo cells (**Suppl. Fig. 4A**).

We next compared α-LGR5 antibody internalisation in cells expressing endogenous levels of LGR5 relative to internalisation of the HER2 cell surface receptor using the well validated, commercial Trastuzumab antibody (α-HER2). For these studies, we used a humanised version of α-LGR5, humanized variant 4 (α-LGR5v4), which retains high-affinity binding to the LGR5 epitope, *Kd* = 2 nM, comparable to the parental murine antibody (**Table 1**). Likewise, α-LGR5v4 detects HEK293T cells overexpressing human and *cynomolgus* LGR5 proteins but not the murine LGR5 (**Suppl. Fig 3**). As a negative control, we used a non-binding version of the antibody generated during the humanisation process, humanized variant 6 (α-LGR5v6), that was non-reactive to the overexpressed LGR5 proteins (**Table 1; Suppl. Fig 3**).

We created fluorescently labelled versions of α-LGR5v4 (Fl-α-LGR5v4) and α-LGR5v6 (Fl-α-LGR5v6) as well as a version of Trastuzumab conjugated to a different fluorophore (Fl-α-HER2) for simultaneous detection. Internalisation of Fl-α-LGR5v4 by LoVo and NALM6 cells was detected by flow cytometry; however, we could only detect Fl-α-HER2 association with LoVo, but not NALM6 cells (**Fig. 4C; Suppl. Fig. 4B**). Significantly less Fl-α-LGR5v6 and the corresponding isotype controls were detected as bound to either LoVo or NALM6 cells (**Fig. 4C; Suppl. Fig. 4B**). Moreover, internalisation of Fl-α-LGR5v4 by either LoVo or NALM6 cells was abrogated by pre-incubation of the antibody with Frag1A (**Suppl. Fig. 4C**).

We next determined the kinetics of endogenous LGR5 internalisation in LoVo cells relative to the HER2 receptor by incubation of Fl-α-LGR5v4 and Fl-α-HER2 for various time periods, up to 180 minutes, and treated the fixed cells with Hoechst and fluorescent phalloidin to visualise nuclei and F-actin. We scored total Fl-α-LGR5v4 and Fl-α-HER2 ‘association’ (bound to the surface or internalised) with cells or ‘internalisation’ by the appearance of Fl-α-LGR5v4 and Fl-α-HER2 puncta inside the cell. Within 5 minutes, both Fl-α-LGR5v4 and Fl-α-HER2 rapidly associated with LoVo cells (**Fig. 4D; Suppl. Fig. 4D**). However, the internalisation kinetics of Fl-α-LGR5v4 and Fl-α-HER2 were different – 100% of the cells associated with Fl-α-LGR5v4 signal show internalisation within 5 minutes whereas only 13% of cells internalised Fl-α-HER2 after 5 minutes, rising to 45% after 180 minutes (**Fig. 4E; Suppl. Fig. 4D**).

Taken together, we have established constitutive and rapid internalisation of LGR5 from a small plasma membrane-localised pool of the protein that can be monitored by bound fluorescent versions of α-LGR5 or α-LGR5v4.

### LGR5 is internalised through the endocytotic pathway

Previous studies with overexpressed LGR5-eGFP have established constitutive internalisation and trafficking to either a LAMP1 positive compartment (26) or the Golgi (29). Internalisation of LGR5 is clathrin and dynamin dependent (21) and is in part regulated by intracellular factors interacting with the C-terminus (29). One study has shown interaction of LGR5 with the cytoskeletal regulator IQGAP1 that in turn influences cortical actin dynamics and cell-cell adhesion (30). However, many of these cellular studies have relied on LGR5 transgene overexpression that may influence internalisation kinetics and cellular distribution. To determine distinguishing features of LGR5 trafficking upon internalisation, we pulsed LoVo and NALM6 cells with Fl-α-LGR5v4 for 60 minutes, fixed and probed the signal from the internalised antibody in relation to molecular probes of specific sub-cellular compartments associated with internalisation and the endocytotic pathway (**Suppl. Fig. 4E**). We generated unbiased co-localisation data sets corresponding to an individual three-dimensional field of view (representative examples of imaging data analysed for LoVo and NALM6 cells are found in **Suppl. Fig. 4F and 4G**, respectively). Image analysis of Fl-α-LGR5v4 and the intracellular markers in LoVo cells (regardless of Fl-α-LGR5v4 internalisation) indicated different degrees of co-localisation, ranging from an average of 22.3% of LGR5 puncta for SNX1 to 8.7% for CD71 (**Fig. 4F**). The level of co-localisation between Fl-α-LGR5v4 and the either GM130 (2.9%) or IQGAP (2.1%) did not significantly deviate from the null hypothesis (no interaction) and were excluded from further analysis.

An inventory of co-incident cellular markers in LoVo cells that have internalised Fl-α-LGR5v4 indicate an even distribution across lysosomes (LAMP1), early endosomes (EEA1 and SNX27), the constitutive internalisation and recycling pathway (CD71) and retromer complex associated vesicles (VPS35 and SNX1), with the majority (28%) associated with SNX1, originally characterised as a marker of vesicles involved in constitutive internalisation of EGFR and its trafficking to lysosomes (31). The least amount of Fl-α-LGR5v4 co-localisation in LoVo cells was with CD71 (Transferrin receptor) a marker of constitutively internalisation and recycling vesicles (**Fig. 4F**) (32).

In NALM6 cells, the distribution of Fl-α-LGR5v4 between the analysed markers was consistent with LoVo cells except for the distribution to SNX1 versus LAMP1 and CD71 which displayed an inverse level of co-localisation; 14% of Fl-α-LGR5v4 co-localised with SNX1 vesicles whereas 16% and 21% co-localised with LAMP1 and CD71 vesicles which suggests a compartmental re-distribution from lysosomal targeting to lysosomal association and plasma membrane recycling in NALM6 cells (**Fig, 4G**). We also observed a small but significant level of co-localisation between Fl-α-LGR5v4 and GM130 (**Fig, 4G**) indicating that LGR5 trafficking to the Golgi network in this cell line may be another distinguishing feature of NALM6 cells.

Taken together, our Fl-α-LGR5v4 internalisation and distribution studies indicate that endogenously expressed LGR5 is rapidly internalised by the endocytotic pathway and is both recycled to the plasma membrane and directed to lysosomal vesicles in both NALM6 and LoVo cells.

### Validation of α-LGR5-based antibody-drug conjugates

Rapid internalisation of α-LGR5 by LoVo and NALM6 cells raises the intriguing prospect of an antibody drug conjugate (ADC)-based therapeutic for targeting cancers that overexpress LGR5. Indeed, two previous studies have used ADC versions of two other LGR5 antibodies for killing LoVo cells *in vitro* and for targeting the LoVo cell tumour model *in vivo* (25,26). We engineered versions of α-LGR5 (murine clone 2), α-LGR5v4 and α-LGR5v6 conjugated to the microtubule poison MMAE. For murine clone 2, we generated both a sulphatase cleavable version (α-LGR5-ADC; (33,34)) and a non-cleavable version (α-LGR5-ADC^NC^) (34). We also generated the IgG1-MMAE conjugate in the cleavable linker format (IgG1-ADC) as a control for our cell killing studies. The α-LGR5-ADC demonstrated almost identical epitope binding affinities as the parental antibody (**Table 1**).

Treatment of NALM6 cells with α-LGR5-ADC for three days lead to effective cell killing, with an EC50 of 4 nM (**Fig. 5A**). α-LGR5-ADC was slightly less effective against the REH pre-B ALL cell line that expresses lower LGR5 levels with an EC50 of 10 nM. We found no effect when treating NALM6 cells with α-LGR5-ADC^NC^ consistent with a previous study that found a non-cleavable version of an LGR5 antibody-based ADC was ineffective at cell killing (26).

**Fig. 5.**
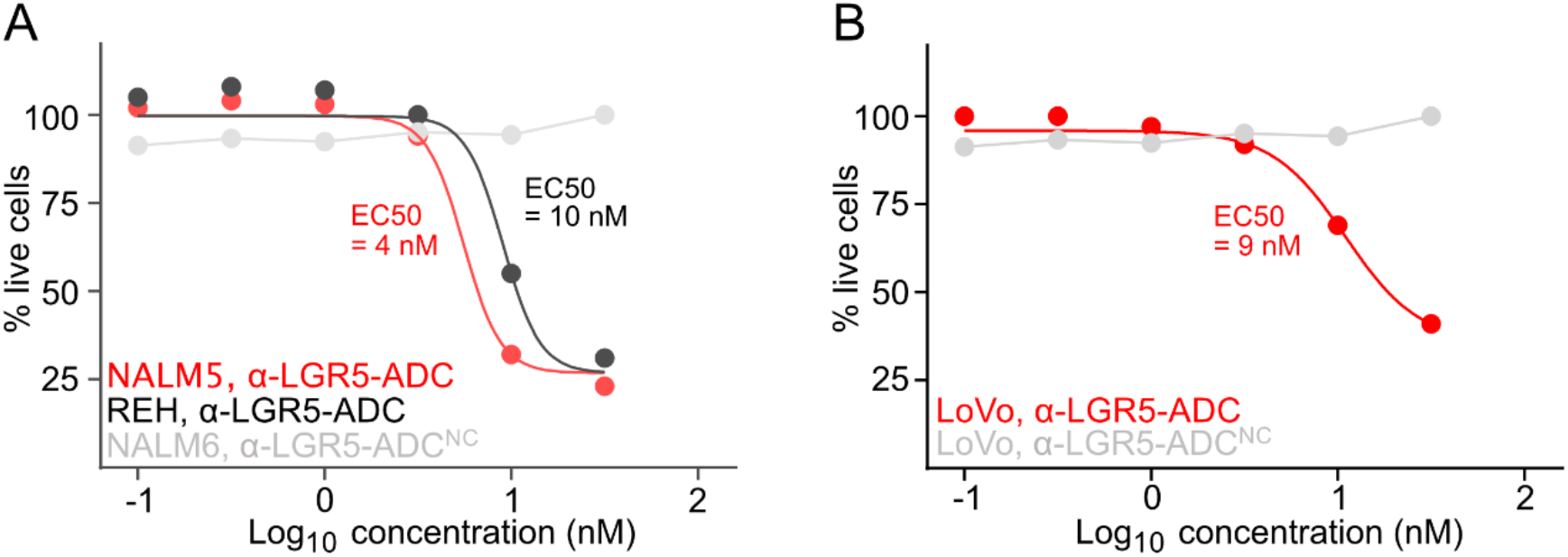
*In vitro* killing of LGR5-expressing cell lines by α-LGR5-ADC. A. Survival of NALM6 or REH cell lines after 72 hours treatment with α-LGR5-ADC. Survival data was fit to a non-linear EC50 shift model yielding EC50 values of 4 and 10 nM, respectively. As control, NALM6 cells were treated with the non-cleavable α-LGR5-ADC^NC^ which did not reduce the cell count after 72 hours. Data is the product of 2 independent experiments for each treatment. B. LoVo cells were treated with either α-LGR5-ADC or the non-cleavable α-LGR5-ADC^NC^. Modelling of cell survival data to a non-linear EC50 shift model yielded an EC50 of 9 nM. Data is the product of 2 independent experiments for each treatment.

We also tested the ability of α-LGR5-ADC to kill LoVo cells, again using α-LGR5-ADC^NC^ as a specificity control. After three days incubation, α-LGR5-ADC effectively killed LoVo cells with an EC50 value of 9 nM (**Fig. 5B**). In contrast, treatment with α-LGR5-ADC^NC^ at concentrations up to 50 nM did not lead to cell death. We conclude that α-LGR5-ADC is highly specific and effectively kills the high LGR5 expressing NALM6 and LoVo cells.

### Targeting NALM6 tumours in vivo with α-LGR5-ADC

To test *in vivo* efficacy of the α-LGR5-ADC we transplanted NALM6 cells stably expressing luciferase into NSG mice. On day 5 post-implantation IVIS imaging was performed and mice were stratified into two groups with equal overall tumour burden. On day 6, 8, 10, and 12 post implantation mice were treated with 5 mg/kg α-LGR5-ADC via tail vein injection (**Fig. 6A**). The control cohort of mice received injections of 5 mg/kg IgG1-ADC control on these days. Tumour burden was monitored at 2–3-day intervals by IVIS imaging. While NALM6 tumours treated with IgG1-ADC grew at a logarithmic rate, we observed tumour regression within 4 days of the initial day 6 of α-LGR5-ADC treatment (**Fig. 6A**). Tumour regression persisted throughout the course of the four treatments after which tumour growth resumed with a 4-day latency period. IVIS imaging on day 18 indicated that the α-LGR5-ADC treatment group had less than 0.5% of the tumour burden of control mice (**Fig 6A, B**). At experimental endpoint on day 20, we also noted a marked reduction in splenic mass (approximately 2-fold) and residual NALM6 cells (approximately 100-fold) as well as absolute numbers of NALM6 cells in the blood (100-fold) and bone marrow (50-fold) of α-LGR5-ADC treated mice (**Fig. 6C**).

**Fig. 6.**
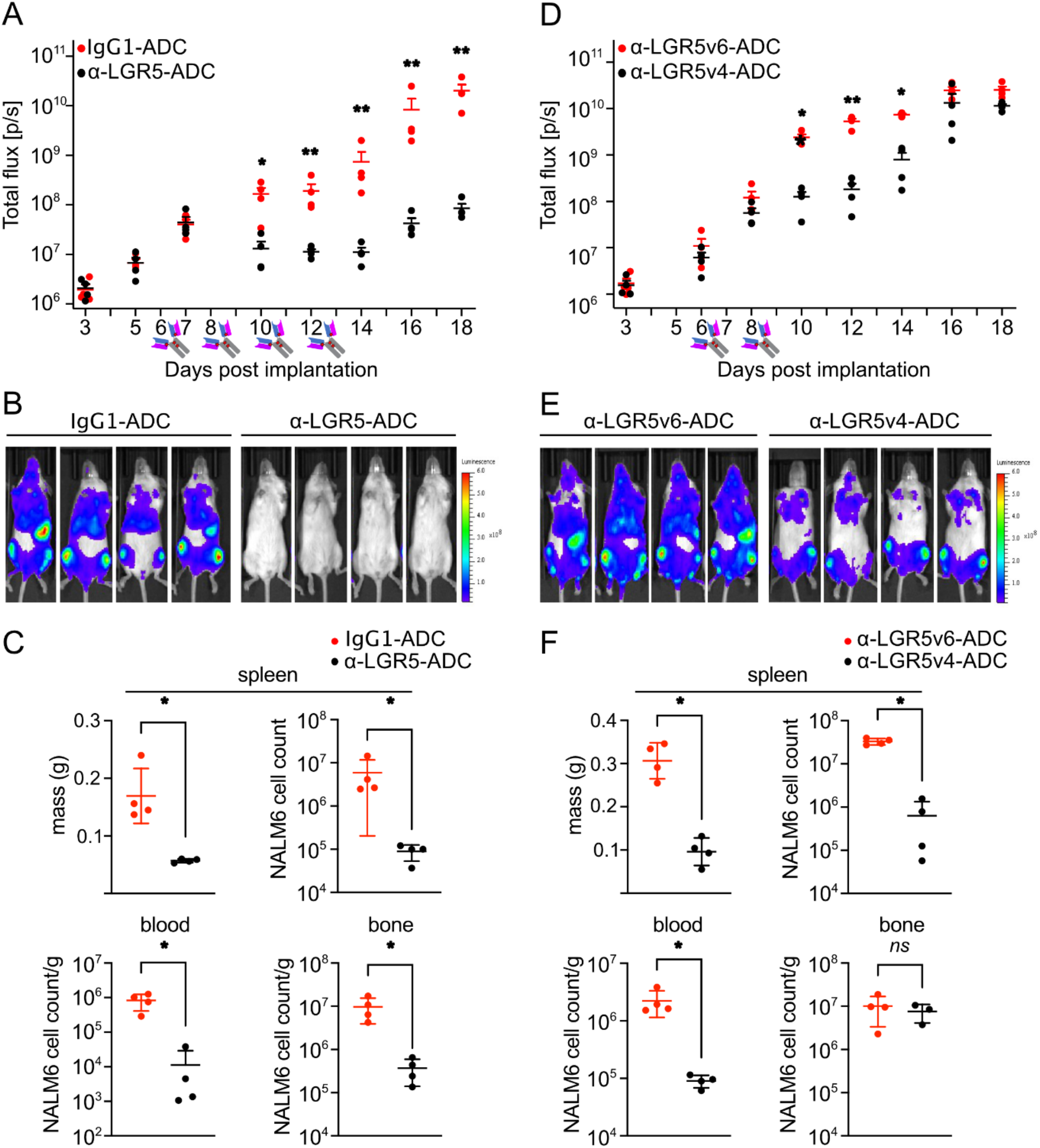
*In vivo* targeting of NALM6 tumours with α-LGR5-ADC. A. Quantification of NALM6 tumour size measured by IVIS imaging over the course of the treatment with either 5 mg/kg α-LGR5-ADC or IgG1-ADC control. Antibody symbols indicate treatments on days 6, 8, 10 and 12 post-implantation. Error bars represent SD at each time point. Significant differences in total flux (bioluminescence), reflective of tumour size, in response to α-LGR5-ADC and IgG1-ADC treatments, was determined by two-tailed t-test and shown at *, *p<0*.*01* and **, *p<0*.*001* levels of significance. B. IVIS images of IgG1-ADC and α-LGR5-ADC treated mice, dorsal view, at experimental endpoint. C. Spleen mass (*top left*) and absolute number of NALM6 cells extracted from spleen (*top right*) of treated mice at experimental endpoint. *Bottom left*, density of NAML6 cells in blood circulation; *bottom right*, NALM6 cells recovered from bone marrow. Error bars represent SD for data from four mice for each treatment. Significant differences were determined by two-tailed t-test at the *, *p<0*.*001* levels of significance. D. Quantification of NALM6 tumour size by IVIS imaging over the course of the treatment with either 5 mg/Kg α-LGR5v4-ADC or α-LGR5v6-ADC. Antibody symbols indicate treatments on days 6 and 8 post-implantation. Error bars represent SD at each timepoint. Significant differences in tumour sizes (total flux) between α-LGR5v4-ADC and α-LGR5v6-ADC treatment groups was determine by two-tailed t-test and shown at *, *p<0*.*01* and **, *p<0*.*001* levels of significance. E. IVIS images of α-LGR5v4-ADC or α-LGR5v6-ADC treated mice, dorsal view, at experimental endpoint. F. Spleen mass (*top left*) and absolute number of NALM6 cells extracted from spleen (*top right*) of treated mice at experimental endpoint. *Bottom left*, density of NALM6 cells in blood circulation; *bottom right*, NALM6 cells recovered from bone marrow. Error bars represent SD for data from four mice for each treatment. Significant differences were determined by two-tailed t-test at the *, *p<0*.*001* levels of significance.

### Targeting NALM6 tumours with a humanized ADC version of α-LGR5

We treated NALM6 tumour bearing mice with only two doses of 5 mg/kg α-LGR5v4-ADC or 5 mg/kg of the control Fl-α-LGR5v6-ADC on days 6 and 8 post implantation. Consistent with our previous *in vivo* trial, tumour growth was largely arrested after a 4-day latency period from the first treatment day. The tumour growth arrest persisted for at least four days after the last α-LGR5v4 treatment after which point tumour growth resumed (**Fig. 6D**). Overall NALM6 tumour burden assessed by IVIS imaging on day 18 for the α-LGR5v4-ADC-treated mice was about half of the control values (**Fig. 6D, E**). As seen with murine α-LGR5-ADC, we noted at experimental endpoint on day 20 a reduction in splenic mass (3-fold) and associated residual NALM6 cell numbers (approximately 50-fold) as well as an approximately 10-fold reduction of NALM6 cells in the blood (**Fig. 6F**). Interestingly, this more modest treatment regime revealed that NALM6 cells were fully retained in the bone marrow where there were no significant differences in total tumour cell number (**Fig. 6F**). Altogether, our *in vivo* trial data indicates that the α-LGR5-ADCs are effective at diminishing tumour growth, but persistent treatment is required for durable responses.

### Application of alternate therapeutic modalities for α-LGR5

We next determined whether α-LGR5 could function in other therapeutic modalities. We generated the humanised α-LGR5 scFv fragment (α-LGR5^scFv^). α-LGR5^scFv^ displays high affinity binding to its LGR5 epitope, with a *Kd* value of 0.77 nM (**Table 1**). We used α-LGR5^scFv^ to generate BiTE constructs and tested the potency of an α-<u>C</u>D3^scFv^-α-<u>L</u>GR5^scFv^ (CL) and α-<u>L</u>GR5^scFv^-α-<u>C</u>D3^scFv^ (LC) BiTE in T cell activation assays. Here we used healthy donor PBMCs as a source of CD4^+^ or CD8^+^ T cells and monitored expression of the T cell activation markers CD25 and CD69 in the presence of tumour cells and respective BiTEs. Low level activation of T cells (less than 3%) was detected for both CD4^+^ and CD8^+^ T cells in the presence of α-LGR5^scFv^. In the absence of target NALM6 cells, LC and CL BiTE treatment also led to low level activation CD4^+^ (8%) and CD8^+^ T cells (5%) (**Figs 7A, B**). However, the presence of both NALM6 target cells, and either BiTE led to 1.5-fold (LC BiTE) and 3-fold (CL BiTE) increase in CD4^+^ T cells activation and a 4-fold (LC BiTE) and 6-fold (CL BiTE) increase in CD8^+^ T cell activation (**Figs 7A, B**). Importantly, we observed effective and specific tumour cell killing when cytotoxic CD8^+^ T cells were directed to NALM6 cells by the respective BiTEs (**Fig. 7C**).

**Fig. 7.**
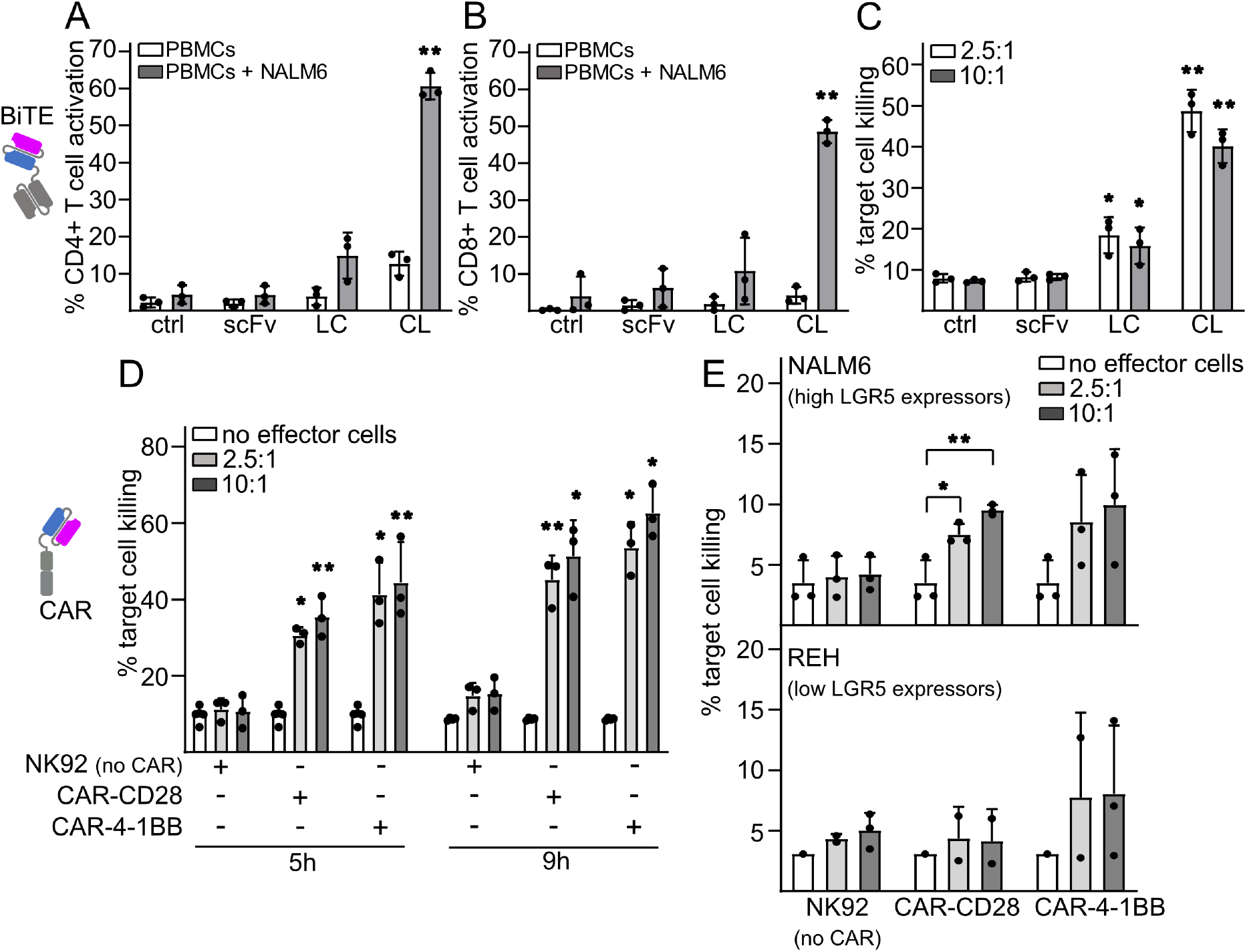
Effective targeting of LGR5-expressing target cells with α-LGR5 BiTEs and CAR NK cells. A. CL- and LC-mediated activation of CD4^+^ T cells in healthy donor PBMCs in the presence or absence of NALM6 target cells determined as percent cells with combined expression of CD25 and CD69. Control (ctrl), no addition of molecule, scFv refers to treatment with the α-LGR5^scFv^ fragment. Error bars represent SD for three independent CD4^+^ T cell activation assays using different healthy donor PBMCs. Significant differences in CD4^+^ activation in between addition of scFv and either the LC or CL BiTE molecules were determined by two-tailed t-test and shown at the **, *p<0*.*001* level of significance. B. CL- and LC-mediated activation of CD8^+^ T cells in healthy donor PBMCs in the presence or absence of NALM6 target cells determined as percent cells with combined expression of CD25 and CD69. scFv is the control α-LGR5^scFv^ fragment. Error bars represent SD for three independent CD8^+^ T cell activation assays using different healthy donor PBMCs. Significant differences in CD8^+^ activation in between addition of scFv and either the LC or CL BiTE molecules were determined by two-tailed t-test and shown at the **, *p<0*.*001* level of significance. C. NALM6 target cell killing by cytotoxic CD8^+^ T cells in the absence (ctrl) or with the addition of scFv or LC or CL BiTEs. Data shown is from three donors and error bars represent SD. The effector to target cell ratios are 10:1 and 5:1. Significant differences in CD8^+^ T cell killing were determined by two-tailed t-test at the *, *p<0*.*005* level of significance. D. Cell killing activity of NK92 cells, or NK92 cells expressing α-LGR5-CAR-CD28- or α-LGR5-CAR-4-1BB. hLGR5-eGFP-overexpressing HEK293T target cells were incubated with effector NK92 cells at effector to target ratios of 2.5:1 and 10:1 for 5h or 9h, respectively. Error bars represent SD for three independent experiments. Significant differences in target cell killing were determined by two-tailed t-test and shown at *, *p<0*.05 and **, *p<0*.*001*. There were no significant differences in target cell killing between cellular ratios of 5:1 or 10:1. E. NALM6 and REH target cell killing activity of NK92 cells or NK92 cells expressing the α-LGR5-CAR-CD28 or the α-LGR5-CAR-4-1BB CARs with effector to target ratios of 2.5:1 and 10:1 after 12 hours incubation (*top* and *bottom panels*, respectively). Error bars represent SD for three independent experiments. Significant differences in target cell killing were determined by two-tailed t-test, at the *, *p<0*.*05* and **, *p<0*.*01*.

Next, we used the α-LGR5^scFv^ to generate second generation CAR constructs for lentiviral delivery, containing the CD28 (α-LGR5-CAR-CD28) or the 4-1BB co-stimulatory module (α-LGR5-CAR-4-1BB), respectively (35). Lentiviruses encoding the CAR construct were used to infect the human Natural Killer cell line NK92, that has recently been suggested as a potential “off-the-shelf” CAR vehicle for therapy (36,37). We tested the ability of NK92 cells expressing either α-LGR5-CAR-CD28 or α-LGR5-CAR-4-1BB to target HEK293T cells expressing LGR5 transgenes. While incubation with NK92 cells alone led to background lysis of approximately 10% in HEK293T cells expressing hLGR5, incubation with NK92 cells expressing the α-LGR5-CAR-CD28 or α-LGR5-CAR-4-1BB transgenes resulted in 35-50% and 50-60% of target cell lysis, respectively (**Fig. 7D**). We observed an approximately 15% increase in cell lysis for NK92 cells expressing the α-LGR5-CARs between 5h and 9h incubations but no differences in HEK293T cell lysis when incubating with cellular ratios of effector cells at 2.5:1 or 10:1. NK92 cells expressing either α-LGR5-CAR-CD28 or α-LGR5-CAR-4-1BB also effectively targeted HEK293T cells expressing the *cynomolgus* LGR5 albeit slightly lower killing was observed. By contrast, no specific lysis was observed for HEK293T cells transfected with mLGR5 (**Suppl. Fig. 6A, B**).

We next evaluated killing potency of NK92 cells expressing α-LGR5-CAR-CD28 or α-LGR5-CAR-4-1BB against NALM6 cells (high endogenous LGR5 expression) and REH cells (low endogenous LGR5 expression). Background killing of target NALM6 cells by NK92 cells was up to 4%, whereas NK92 cells expressing α-LGR5-CAR-CD28 killed 7.5% and 9.5% at effector target cell ratios of 2.5:1 and 10:1, respectively (**Fig. 7E**). NK92 cells expressing α-LGR5-CAR-4-1BB also showed increased cell killing relative to NK92 cells however, the difference was not significant. We also observed an increase, albeit not significant, in killing of REH cells between NK92 cells and NK cells expressing either α-LGR5-CAR-CD28 or α-LGR5-CAR-4-1BB. (**Suppl. Fig. 6C**).

Taken together, we have established high *in vitro* specificity and efficacy for the α-LGR5 scFv fragment in the BiTE and CAR-NK modalities.

## DISCUSSION

LGR5 is an established stem cell marker in a number of murine tissues (38–40) and studies using genetically engineered mouse models (GEMMs) and human CRC organoids have ascribed LGR5 a functional role in maintaining the proliferative compartment in tumours (15,25,26). Here we report the development, validation and characterisation of novel antibodies raised against the extracellular domain of LGR5 with clear line of sight to therapeutic application. α-LGR5 is unique from previously reported α-LGR5 antibodies (25,26) in its CDR3 regions and its target epitope on the LGR5 protein, localised to the N-terminal 15 amino acids of the extracellular domain. The parental murine α-LGR5 antibodies, its humanised and ADC versions and the scFv fragment are all high affinity binders compared to existing LGR5/hybrid therapeutic antibody candidates in development (**Table 2**).

**Table 2.**
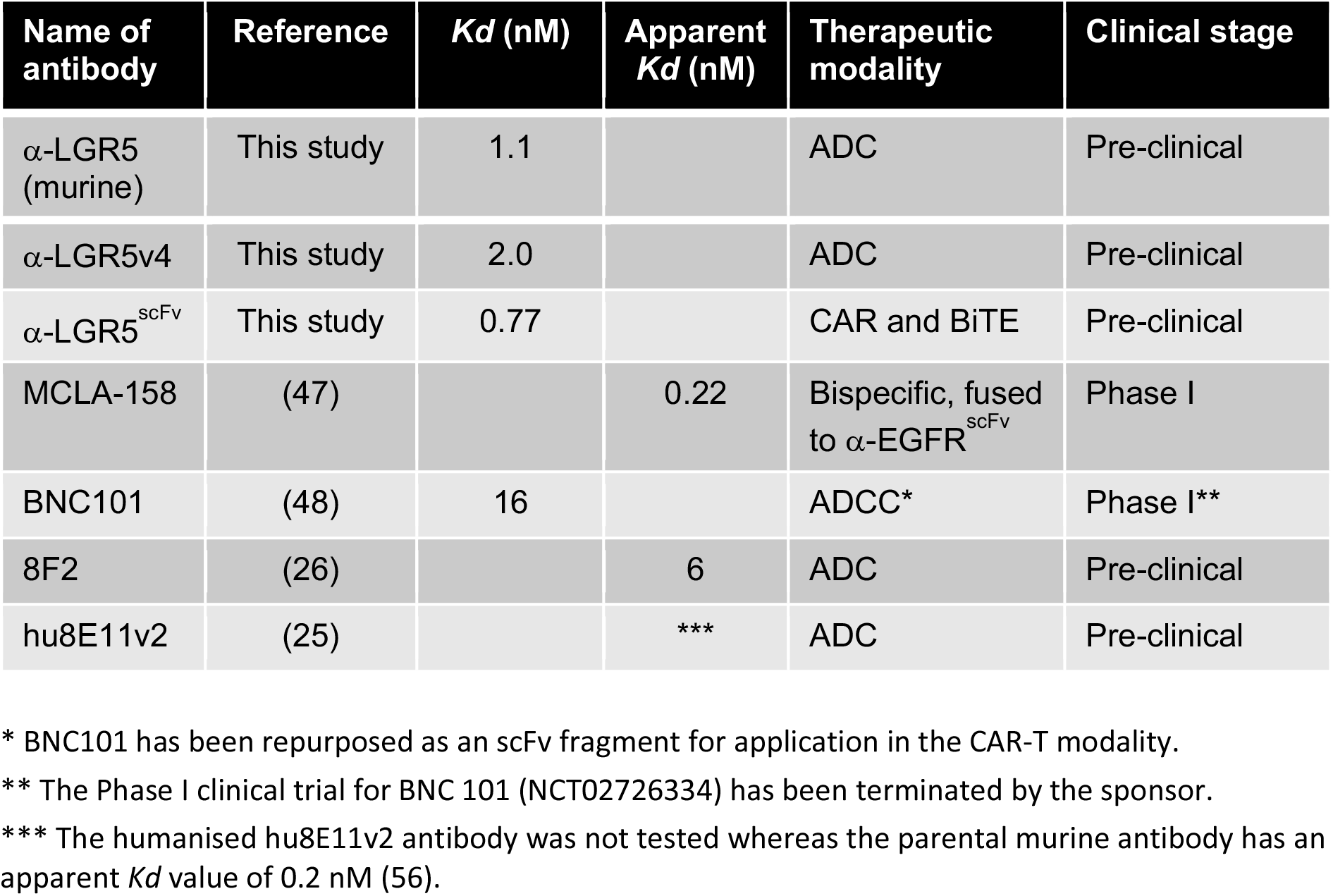
Binding affinities for therapeutic LGR5 antibodies.

| Name of antibody | Reference | *K<sub>d</sub>* (nM) | Apparent *K<sub>d</sub>* (nM) | Therapeutic modality | Clinical stage |
| --- | --- | --- | --- | --- | --- |
| α-LGR5 (murine) | This study | 1.1 |  | ADC | Pre-clinical |
| α-LGR5v4 | This study | 2.0 |  | ADC | Pre-clinical |
| α-LGR5<sup>scFv</sup> | This study | 0.77 |  | CAR and BiTE | Pre-clinical |
| MCLA-158 | (47) |  | 0.22 | Bispecific, fused to α-EGFR<sup>scFv</sup> | Phase I |
| BNC101 | (48) | 16 |  | ADCC* | Phase I** |
| 8F2 | (26) |  | 6 | ADC | Pre-clinical |
| hu8E11v2 | (25) |  | *** | ADC | Pre-clinical |

\* BNC101 has been repurposed as an scFv fragment for application in the CAR-T modality.
\*\* The Phase I clinical trial for BNC 101 (NCT02726334) has been terminated by the sponsor.
\*\*\* The humanised hu8E11v2 antibody was not tested whereas the parental murine antibody has an apparent *K<sub>d</sub>* value of 0.2 nM (56).

**Table 3.** Antibodies and probes used in the study.

| Antibody or Probe | Clone | Conjugate | Application | Manufacturer |
| --- | --- | --- | --- | --- |
| **Primary antibodies** |  |  |  |  |
| α-vinculin | 4650 | none | WB | Cell Signalling Technology |
| α-β-actin | AC-15 | none | WB | Sigma-Aldrich |
| α-HER2 | Trastuzumab | none | IF and ADC | Roche |
| α-CD71 | OKT9 | none | IF | Thermo Fisher Scientific |
| α-SNX1 | 51/SNX1 | none | IF | BD Biosciences |
| α-SNX27 | 1C6 | none | IF | Abcam |
| α-EEA1 | 14/EEA1 | none | IF | BD Biosciences |
| α-VPS35 | Goat polyclonal | none | IF | Novus biologicals |
| α-IQGAP1 | D6E3J | none | IF | Cell Signalling Technology |
| α-LAMP1 | H4A3 | none | IF | Developmental Studies Hybridoma Bank |
| α-GM130 | 35/GM130 | none | IF | BD Biosciences |
| α-HA | 12CA5 | none | WB | a kind gift from Sean Munroe |
| α-B220 | RA3-6B2 | BV785 | FACS | Biolegend |
| α-CD3 | UCHT1 | BUV395 | FACS | BD Biosciences |
| α-CD3 | UCHT1 | BV510 | FACS | Biolegend |
| α-CD4 | RPA-T4 | BV605 | FACS | Biolegend |
| α-CD8 | RPA-T8 | BV711 | FACS | Biolegend |
| α-CD8 | RPA-T8 | BV737 | FACS | Biolegend |
| α-CD19 | HIB19 | BV421 | FACS | Biolegend |
| α-CD19 | SJ25C1 | BUV395 | FACS | BD Biosciences |
| α-CD34 | 581 | PE | FACS | Biolegend |
| α-CD45 | HI30 | BUV737 | FACS | BD Biosciences |
| α-CD45 | HI30 | FITC | FACS | Biolegend |
| α-CD95 | DX2 | Bv421 | FACS | Biolegend |
| α-CD127 | A019D5 | BV711 | FACS | Biolegend |
| α-CCR7 | G043H7 | PE-Cy7 | FACS | Biolegend |
| α-HLA-DR | L243 | BV510 | FACS | Biolegend |
| α-IgM | HI100 | PE-Cy7 | FACS | Biolegend |
| α-CD45.1 | A20 | FITC | FACS | Biolegend |
| α-CD25 | M-A251 | BV421 | FACS | BD Biosciences |
| α-CD69 | FN50 | APC, FN50 | FACS | Biolegend |
| α-FLAG | M2 | Agarose beads | WB | Sigma-Aldrich |
| **Secondary antibodies** |  |  |  |  |
| Goat-α-mouse IgG | | HRP | WB | Dako |
| Goat-α-rabbit IgG | | HRP | WB | Dako |
| Donkey-α-mouse IgG (H+L) Highly Cross-Absorbed | | AF488 | IF | Thermo Fisher Scientific |
| Goat-α-rabbit IgG (H+L) Highly Cross-Absorbed | | AF488 | IF | Thermo Fisher Scientific |
| Donkey-α-goat IgG (H+L) Highly Cross-Absorbed | | AF488 | IF | Thermo Fisher Scientific |
| AffiniPure Donkey-α-human IgG (H+L) | | AF647 | IF | Jackson ImmunoResearch |
| **Probes** |  |  |  |  |
| Phalloidin |  | AF488, AF568, AF647 | IF | Life Technology |
| Hoechst 33342 |  | n/a | IF | Thermo Fisher Scientific |

α-LGR5 is a highly versatile research tool compatible with a range of techniques such as western blot, immunofluorescence and flow cytometry and has proven to be a sensitive and specific reagent for determining cellular levels and localisation of LGR5 protein in healthy and malignant tissue. Indeed, we were able to conduct extended analysis of LGR5 protein expression, an important extension to previous transcriptomic analyses (25,26) (as well as the present study), comparing normal and cancer tissues. We note the prominent differences between the results obtained from our and previous transcript analyses of LGR5 and actual LGR5 protein expression exemplified in brain and ovarian cancers. While both malignancies consistently score high for LGR5 transcript levels, we only detected low LGR5 protein levels in few cells, in less than 10% of the tumours. The disparity may be due to the ability to distinguish LGR5 expression in epithelial tissue by immune fluorescence versus bulk transcript levels in biopsies or the use of different tissue collection protocols amongst sample cohorts and highlights the need for serial comparison of LGR5 protein levels as a true indicator of tissue and tumour expression.

We have established elevated LGR5 expression as a characteristic of CRC, HCC and pre-B ALL. Importantly, our tissue census, both transcriptional and at the level of LGR5 protein, indicates that normal tissues harbour very low to undetectable LGR5 levels, paving the way for therapeutic targeting of malignancies that overexpress the protein. Indeed, our approach has established CRC, HCC and pre-B ALL as priority cancer targets for α-LGR5-based therapeutics and enables future studies to determine other targetable cancer types and stratifying LGR5 overexpression as a prognostic marker. This is particularly relevant to the assessment of LGR5 protein levels in HCC – high LGR5 protein expression may be used to further stratify the HCC subset with activating mutations in β-catenin, which are characterised by low T cell infiltration and thus referred to as immune deserts (41,42). The prediction is that this HCC subset will be refractory to both checkpoint inhibition and cellular therapies. Indeed, recent reporting from the CheckMate 459 trial (NCT02576509) evaluating nivolumab (PD1 checkpoint inhibitor) versus sorafenib (small molecule kinase inhibitor) in HCC failed to meet its endpoint target of improved overall survival (41,43). However, we propose that exclusion of the high LGR5 expressing patient cohort may prove to be important in the outcome of the trial. Hence, stratification of HCC patients with high levels of LGR5 using α-LGR5 presents an intriguing biomarker opportunity that both reports the β-catenin mutant subset and supports the use of therapeutic molecules such as α-LGR5-ADC, where drug efficacy is not dependent on immune infiltration.

The range of techniques that are compatible with α-LGR5 allowed us to establish the steady-state cellular distribution of the LGR5 protein - low level expression on the cell surface with the majority associated with intracellular puncta due to rapid constitutive internalisation – within 5 minutes LGR5 reaches its maximal level of internalization. By contrast, HER2 requires 180 minutes to reach a maximal internalisation of 45%. The concern with long retention times on the cell surface is the increased possibility of ADC linker hydrolysis resulting in off-target bystander toxicity.

In addition to its utility as a research tool, we demonstrate therapeutic application of α-LGR5 as an ADC, BiTE and CAR. The highly dynamic nature of LGR5 trafficking, the window of LGR5 expression between healthy cells and specific cancers, and the ability of the cell surface pool of the protein to internalise cargo in overexpressing cells supports α-LGR5 as an excellent platform for ADC-based therapeutics. Indeed, α-LGR5-ADCs have been previously road-tested for targeting LoVo cell tumours (25,26). The present study expands the application of α-LGR5-ADC by targeting NALM6 pre-B ALL tumours. Our use of two different α-LGR5-ADC treatment regimens lower than reported by previous studies has revealed important pharmacodynamic properties of this therapeutic strategy for targeting blood malignancies: the reduction in NALM6 tumour size manifests up to 4 days post treatment and is only sustained for up to 4 days post-treatment at which point we observe tumour re-growth. Our post-tissue census of residual tumour cells at experimental endpoint indicates that tumour relapse may stem from a reservoir of NALM6 cells within the bone marrow and this tumour compartment can only be effectively targeted by four or more rounds of our α-LGR5-ADC treatment regime. These data are informative for clinical application of α-LGR5-ADC in targeting blood malignancies and indicate that effective targeting requires increased amounts of α-LGR5-ADC with sustained rounds of treatment.

We did not observe off-target toxicities in α-LGR5-ADC treated mice; however, α-LGR5 does not cross react with the murine LGR5 and it will be important to evaluate on-target, off-tumour toxicities in an appropriate model. Previous trials of α-LGR5-ADCs targeting the LoVo tumour model that utilised a murine cross-reactive antibody were conducted at greater levels of drug or with prophylactic treatment and observed no associated treatment toxicities (25,26).

We show that α-LGR5 is a particularly versatile therapeutic antibody and the corresponding scFv fragment is effective in both BiTE and CAR modalities. BiTEs in the form of α-LGR5^scFv^-α-CD3ε^scFv^ fusions displayed high specificity in the activation of CD4^+^ and CD8^+^ T cells and effectively promoted CD8^+^ T cell-mediated NALM6 tumour cell destruction. Moreover, the two second-generation CAR molecules we developed and expressed in NK92 cells effectively lysed human and *cynomolgus* LGR5-overexpressing HEK293T cells but not cells overexpressing the murine LGR5. Specificity of the NK92-CAR was also apparent with more effective targeting of NALM6 cells relative to REH cells which express lower amounts of the LGR5 protein.

Taken together, our study has important implications for cancer research and therapeutics: (1) the highly specific α-LGR5 antibody we have developed will be particularly useful as a research tool for determining novel cell biology of human LGR5, (2) LGR5-expressing tumour cells have been validated as *bone fide* therapeutic targets in CRC, HCC and pre-B ALL with the intriguing possibility of identifying further LGR5 expressing cancers types;, and (3) the demonstration that α-LGR5 is an adaptable therapeutic antibody for targeting cancer cells in three different therapeutic modalities and can thus accommodate different pharmacodynamic requirements of various LGR5^+^ tumour types.

## MATERIALS AND METHODS

### Plasmid constructs

The plasmid for hLGR5-eGFP expression plasmid has been previously described (29). All other LGR transgenes used in the study were constructed by direct replacement of the LGR5 coding sequence with PCR amplicons from the corresponding LGR-family coding sequence by either Gibson assembly (New England Biolabs) or restriction enzyme cloning. LGR-family coding sequences were sourced as follows: hLGR4 (HG15689), Sino Biological; mLGR4 (MR219497) Origene; mLGR5 (MR219702) Origene; and hLGR6(LGR6_OHu16329D) GenScript.

Sequencing of antibody hybridoma clones and generation of plasmids encoding heavy and light chains of the humanised α-LGR5 antibodies (α-LGR5v4 and α-LGR5v6) and the scFv fragment were obtained from Absolute Antibody.

### Antibodies and probes

Antibodies and probes used in the study are listed in **Suppl. Table 1**.

### Generation of α-LGR5 antibodies

The fusion was made from splenocytes of NMRI mice or SPRD rats (Taconic), which were SC immunized twice at a 14-day interval with 30 mg of the LGR5 N-terminal fragment glutaraldehyde coupled to diphtheria toxoid. The antigen was administered with the GERBU Pä adjuvant according to the manufacturer’s recommendations. Four days prior to the fusion, the animals received an IV injection boost of 15 mg antigen administered with adrenaline. Fusions and screenings were performed with the mouse SP2 myeloma cell line as the fusion partner by established methods.

Purification of all immunoglobin fractions was carried out by absorption of either 1 L of hybridoma supernatants, or 200 ml of conditioned media from heavy and light chain expressing HEK293T cells, to 3 ml packed volume of Protein G fast flow Sepharose equilibrated in PBS. After extensive washing of the columns with PBS, the immunoglobin fraction was eluted with 100 mM glycine, pH 2.7 and immediately neutralized in 200 mM Tris pH 8.0.

### Preparation of proteins for bacterial expression

The coding sequence for the N-terminal 100 amino acids of human LGR5 lacking the signal peptide was inserted into the pGEX-4T1 bacterial expression vector and expressed in E. coli XL1 blue. The expressed GST-fusion protein was absorbed to glutathione-Sepharose 4B (Sigma-Aldrich) followed by extensive washing with phosphate buffered saline (PBS) and elution from the matrix with 20 mM glutathione. The eluate was digested overnight with 2 U of thrombin protease (Sigma) and the LGR5 N-terminal fragment used as the antigen for α-LGR5 antibody generation was resolved on a Superdex 75 10/300 gel filtration column (Cytiva) equilibrated in PBS using the ÄKTA pure system.

Bacterial expression plasmids for RAD display of LGR5 N-terminal fragments were generated by Gibson assembly, expressed in bacteria and purified by heat denaturation and Ni-Sepharose affinity chromatography as previously described (22).

### Mammalian cell lines

Cell lines were purchased from the European Collection of Cell Cultures and have been authenticated by short tandem repeat (STR) DNA profiling. Upon receipt, cell lines were frozen, and individual aliquots were taken into culture, typically for analysis within <10 passages. Cells were grown in a humidified incubator at 37^∘^C and 5% CO_2_ and tested mycoplasma negative (MycoProbe^®^ Mycoplasma Detection Kit, R&D systems). HEK293T cells, and the colorectal cancer cell lines LoVo, SW480, HT29, HCT116, CaCo and DLD1, were maintained in DMEM medium supplemented with 10% heat-inactivated FCS (Gibco) and 100 U/ml penicillin/streptomycin (Gibco). pre-B ALL cell lines NALM6, REH, 697, RS4-11, HAL-01, NALM16, SupB15 KOPN8, and MHH-CALL2 were maintained in RPMI-1640 (Gibco) medium supplemented with 10% heat-inactivated FCS (Gibco) and 100 U/ml penicillin/streptomycin (Gibco).

### Cellular assays and manipulation

Indirect immunofluorescence, western blot analysis, flow cytometry and TopFlash assays have been previously described (44,45). For immunofluorescence, detection was carried out by confocal spinning disc microscopy using an Andor Dragonfly 500 (Oxford Instruments). Images were processed using Imaris software (Bitplane/Oxford Instruments). Flow cytometry was carried out with a BD LSR Fortessa or BD LSR Symphony cell analyzer using the BD FACSDiva software (BD Biosciences Inc.). Quantitative real-time PCR (qRT-PCR) has been previously described (45) and used Taqman probes specific for human LGR5 (Life Technology, Hs00969422_m1) and for TBP (Life Technology, Hs00427620_m1) as control housekeeping gene. The Prism software package was used to graph data sets from TopFlash assays and qRT-PCR experiments and for statistical analysis using two-tailed Students t-test.

For overexpression of LGR family proteins, HEK 293T cells were transfected with the corresponding plasmids using lipofectamine 2000 (Life Technology) according to manufacturer’s recommendations. Cells were transfected overnight and recovered in culture media for an additional 16 hours prior to immunofluorescence, western blot or flow cytometry.

Generation of α-LGR5v4 and α-LGR5v6 was carried out by transfection of HEK293T 4 x T175 flasks with 15 μg of each encoding plasmid (Absolute Antibody). Media containing antibodies was collected at 2- and 4-days post-transfection and subject to Protein G purification. The fluorescent versions of α-LGR5 (Fl-α-LGR5), α-LGR5v4 (Fl-α-LGR5v4) and α-LGR5v6 (Fl-α-LGR5v6) were generated using the Dylight TM650 Antibody labelling kit (Thermo Fisher Scientific). Fluorescent Trastuzumab (Fl-α-HER2) was generated using the Dylight TM550 Antibody labelling kit, respectively. In some controls for flow cytometry and immunofluorescence, Fl-α-LGR5 was pre-incubated with supra-stoichiometric amounts of RAD-displayed Frag1A or Frag1B, or the Frag1A peptide (Cambridge Peptides) at a molar ratio of 10:1.

Antibody internalisation assays were carried out by incubation of cells with 20 μg/ml of Fl-α-LGR5 or Fl-α-HER2 for various timepoints, followed by fixation in 4% paraformaldehyde and immunofluorescence. Images derived from z-stacks of cells were analysed for Fl-α-LGR5v4 and fluorescent signals from antibodies against various cellular markers, fluorescent phalloidin to visualise cortical actin, and Hoechst 33342 to visualise nuclei. Images were processed and analysed used Arivis Vision 4D software. Image segmentation to delineate fluorescent features: whole cells, delineated by cortical F-actin, nuclei and puncta) in 3D was carried out using Blob Finder. The arivis Vision 4D software was then used to classify puncta associated with the cell membrane or within the cells and determine the degree of co-localisation between LGR5 and compartment-specific markers – co-localisation is defined as >50% overlap with LGR5 puncta.

For the internalisation kinetics experiments, images were analysed using arivis Vision 4D software, combined with a deep-learning segmentation using Cellpose. Two regions were defined: cell outer membrane, and cell cytoplasm. Segmented dotted signal corresponding to LGR5, as well as segments-like signal corresponding to Trastuzumab, were classified according to their location within these two regions. Ratiometric analysis was performed to quantify the internalisation of both markers at the various time points.

### TCGA data mining for LGR5 expression in cancers

Publicly available gene expression data (RNAseq V2) from The Cancer Genome Atlas (TCGA; https://www.cancer.gov/about-nci/organization/ccg/research/structural-genomics/tcga) were downloaded using Firebrowse (http://firebrowse.org/). Gene-level read counts were quantile normalised using Voom (Law et al., 2014) and (log2 median-centred) *LGR5* gene expression was determined for each sample. Tumour subtypes for which more than 70% of samples had higher than pan-cancer median LGR5 expression were triaged as “high LGR5 tumours”.

### Patient samples and immunofluorescence detection of LGR5 and β-catenin

All human tissue biopsies used in the study were paraformaldehyde-fixed paraffin embedded and probed with α-LGR5 and an antibody to β-catenin and visualised by fluorescent secondary antibodies to mouse labelled with Alexa 488 and rabbit labelled with Alexa 555. All immune stained samples were imaged using the PhenoImager HT™ Automated Quantitative Pathology Imaging System (Akoya Biosciences). Scoring of all human biopsies for simultaneous LGR5 and β-catenin expression was performed by an individual blind to the provenance of the samples and graded from no expression – 0 to high expression – 3 for all sample sets. The Prism software package was used for plotting of LGR5 or β-catenin expression levels for all biopsy sample sets and for determining statistical differences using two-tailed students t-test. Unless otherwise notes, all relevant legal and ethical guidelines of the Addenbrooke’s Hospital (Cambridge, UK) were followed for collection of samples and provision for the present study. Informed consent for research application was obtained from all subjects.

LGR5 expression in individual colorectal cancer cases and adjacent healthy tissue were determined for biopsies provided by Dr Olivier Giger (OG; (IRAS: 162057). Within these tissue samples, regions were annotated as normal, dysplastic or invasive tissue from consecutive H&E sections. LGR5 and β-catenin protein levels were additionally determined in CRC by immunofluorescence using the Bern CRC sample set, provided by Dr Inti Zlobec. The Bern CRC sample set is a highly annotated tumour microarray (TMA) consisting of 160 individual cases in duplicate with determined phenotypic feature - gender, age, tumour stage, therapeutic intervention, and MSI status. Biopsies used in the construction of the TMA were collected under ethics 2020-00498 granted by the Ethical Committee of the Canton of Bern, Switzerland. All relevant guidelines of the Institute of Pathology, University of Bern, Canton of Bern, Switzerland were followed for construction of the TMA.

The Cambridge HCC TMA consists of 104 human liver samples and was collected by Drs Sarah Aitken and Matthew Hoare with informed consent from Addenbrooke’s Hospital, Cambridge, UK, according to procedures approved by the East of England Local Research Ethics Committee (16/NI/0196 and 20/EE/0109). Liver samples classified as healthy were obtained from resections from females with inflammatory adenoma of the liver (2 individuals) or focal nodular hyperplasia (2 individuals) or from a male with an HNF1α-inactivated adenoma. All biopsies of healthy liver tissue were taken from patients between the ages of 25 and 36.

High grade serous ovarian carcinoma (HGSOC) samples comprising the Cambridge ovarian cancer TMA were provided by Prof James Brenton. Tumour samples were obtained from patients enrolled in the Cambridge Translational Cancer Research Ovarian Study 04 (CTCROV04, short OV04) study approved by the Institutional Ethics Committee (REC08/H0306/61). Samples were processed following standardised operating protocols as outlined in the OV04 study design. Tissue quality was assessed using haematoxylin and eosin (H&E) sections, and high purity regions were selected for tissue microarray (TMA) generation (using 0.1 cm cores). The TMA consisted of healthy fallopian tube (FT; 27 samples), 28 ovarian cancer cases (OvC) and 14 omentum cancer cases (OmC).

The Cambridge brain cancer TMA consists of 5 samples of healthy brain tissue, 5 from low grade glioma and 5 from glioblastoma that were collected via the ICARUS biorepository, Addenbrooke’s Hospital, according to approved local research ethics (LREC 18/EE/0172).

Sections of PDAC and healthy pancreas were provided by Dr Eva Serrao and were obtained from the Cambridge University Hospital Human Tissue bank, Cambridge, UK, according to procedures approved by the Cambridge South Research Ethics Committee (REC18/EE/0227).

Primary haematological malignancy samples used in this study were provided by Blood Cancer UK Childhood Leukaemia Cell Bank (LREC 16/SW/0219) and Cambridge Blood and Stem Cell Biobank (LREC 18/EE/0199) as well as the Blood Cancer UK Biobank (REC16/SW/0219) according to procedures approved by the South West-Central Bristol Research Ethics Committee.

Buffy Coats from healthy donors were acquired from NHS Blood and Transplant (Cambridge) under appropriate ethics (Research into Altered Lymphocyte Function in Health and Disease, REC reference: 17/YH/0304). PBMCs were isolated using SepMate PBMC Isolation Tubes (Stemcell Technologies) and B cells and CD8^+^ T cells were isolated from these using the human CD19 Microbeads (Miltenyi Biotec) or the human CD8^+^ isolation kit (Miltenyi Biotec).

### ADC generation and *in vitro* killing assays

All antibodies and IgG1 (Sigma-Aldrich) were coupled to MMAE through a divinyl pyrimidine bridging linker inserted within the heavy chain-light chain disulphide linkages for precise drug-to-antibody ratio of 4 (34). For *in vitro* killing assays, LoVo target cells were seeded into opaque 96-well plate for overnight, allowing settlement before treatment using ADCs with cleavable linker or non-cleavable control linker at doses of 30, 10, 3, 1, 0.3, and 0.1 nM for 3 days. NALM6 and REH target cells were seeded into opaque 96-well plate and treated directly by ADCs with cleavable linker or non-cleavable control linker at doses of 30, 10, 3, 1, 0.3, and 0.1 nM for 3 days. On day 3, cell viability was evaluated by CellTiter-Glo 2.0 Cell Viability Assay (Promega) according to manufacturer’s instructions. Bioluminescence was measured using a CLARIOStar (BMG Labtech).

### *In vivo* therapeutic efficacy of mouse and humanised ADCs

NSG mice (NOD scid gamma; strain *NOD*.*Cg-Prkdc*^*scid*^ *Il2rg*^*tm1Wjl*^*/SzJ*) were purchased from Charles River UK Ltd (Margate, UK) and housed under specific pathogen-free conditions at the University of Cambridge, CRUK Cambridge Institute in accordance with UK Home Office regulations. NALM6 cells were transduced with lentivirus encoding a LucEYFP reporter (the plasmid was a kind gift of Kevin Brindle).

Immunodeficient NSG mice were injected intravenously with LucEYFP-expressing NALM6 cells by tail vein injection and monitored for weight loss and IVIS imaging in a 2–3-day interval. For IVIS imaging, mice were given D-luciferin at a dose of 150 mg/kg by i.p. 10 minutes prior to imaging by IVIS2000 (PerkinElmer) under general anaesthesia with isoflurane. Mice were randomised into control and treatment groups according to the bioluminescence signal detected on day 5 after xenograft.

Treatment with mIgG-ADC or α-LGR5-ADC at a dose of 5 mg/kg was carried out by i.v. injection starting on day 6 for 4 times every other day. Treatment with α-LGR5v6-ADC (control) or α-LGR5v4-ADC at dose of 5 mg/kg was carried out by i.v. injection on days six and eight. At the experimental endpoint on day 20, all animals were euthanised in accordance with Schedule 1 of the Animals (Scientific Procedures) Act 1986. Spleen, blood, heart, kidney, lung, liver, small intestine, femur and tibia were collected at the experimental endpoint for histology and/or FACS.

Single cell suspensions prepared from spleen, blood, and bone marrow of femur and tibia were stained with eflour780 fixable live/dead, followed by fluorescence-conjugated antibody against human CD19 for identification of NALM6 cells (CD19^+^ EYFP^+^) by FACS analysis. Absolute number of NALM6 cells was determined using AccuCheck Counting Beads (Thermo-Fisher Scientific).

### BiTE production, T cell activation and killing assay

Expression plasmids containing the coding region for transgenic BiTE versions of α-LGR5^scFv^ fused to the scFv antibody fragment of CD3ε were constructed in both orientations: either with LGR5^scFv^ at the N-terminus, LGR5^scFv^-CD3ε^scFv^ (LC BiTE), or with CD3ε^scFv^ at the N-terminus (CL BiTE). The CL and LC BITEs were engineered to include an N-terminal signal sequence and FLAG epitope tag. Both BiTEs were purified from conditioned media of transfected HEK293T cells using HiTrap Protein L column (Cytiva) chromatography.

T cell activation assays were initiated by adding 3.7 nM CL or LC BiTEs or α-LGR5^scFv^ as a negative control to a mixture of 10^6^ PBMCs and 10^6^ NALM6 cells. After 24-hour incubation, cells were stained with eflour780 fixable live/dead dye, followed by fluorescence-conjugated antibodies against CD4, CD8, CD25 and CD69. Flow cytometry was used to assess expression of the T cell activation markers CD25 and CD69 on CD4^+^ and CD8^+^ T cells.

BiTE-mediated killing assays were carried out using cytotoxic CD8^+^ T cells. For this, CD8^+^ T cells were isolated from PBMCs as described and stimulated for 72h with 25µL/mL ImmunoCult™ Human CD3/CD28/CD2 T Cell Activator (STEMCELL). CD8^+^ T cells were cultured in TexMACS media (Miltenyi Biotec) supplemented with 100 U/ml human IL-2 (Miltenyi Biotec) and 100 U/ml Penicillin/Streptomycin (Gibco). Cells were restimulated at 1×10^6^ cells/mL concentration for 48 h on day 8-10 with 1 µg/mL plate-bound anti-CD3ε antibody (clone UCHT1, Biolegend). BiTE-mediated killing assays were carried out on day 14-16 of culture.

NALM6 target cells were labelled with CellVue membrane dye (CellVue Claret Far Red Fluorescent Cell Linker Kit protocol; Sigma-Aldrich) and 6.25 × 10^4^ or 1.25 × 10^5^ cytotoxic CD8^+^ T cells were used at indicated effector to target rations. After 6 hours of co-culture at 37°C, cells were labelled with eFluor780 fixable live/dead dye (ThermoFisher) and NALM6 cells were analysed for viability by flow cytometry.

### CAR-NK cell production and killing assay

The α-CD19-CAR lentiviral plasmid was a kind gift from Dr John James (University of Warwick) (46) and contained the coding region for the α-CD19 scFv domain (clone FM363) fused to the CD8 stalk and transmembrane regions, a 4-1BB or CD28 intracellular costimulatory domain, respectively, and mScarlet.

NALM6, REH or HEK293T cells overexpressing hLGR5-eGFP, mLGR5-eGFP or cLGR5-eGFP were used as target cells in the killing assay and were pre-loaded with CellVue membrane dye (CellVue Claret Far Red Fluorescent Cell Linker Kit protocol; Sigma-Aldrich) and 2.5 × 10^6^ or 6.25 × 10^5^ NK92 cells or CAR expressing NK92 cells were used at indicated effector to target ratios. At designated timepoints (5, 6, 9 or 12 hours), percent target cell killing was assayed by flow cytometry.

## List of Supplementary Materials

Fig. S1. Specificity of LGR5 antibodies generated in the study.

Fig. S2. Supplemental Figure 2. LGR5 expression analysis in cancers

Fig. S3. Validation of α-LGR5v4

Fig. S4. LGR5 internalisation in NALM6 and LoVo cells.

Fig. S5. α-LGR5-mediated cell killing in BiTE and CAR modalities

## Acknowledgements

We acknowledge the contribution of the CRUK CI Core Facilities: the BRU facility for excellent animal husbandry; the Preclinical Imaging Core for help with *in vivo* imaging (IVIS), the Light Microscopy and the Histopathology core, as well as the Compliance and Biobanking and Flow Cytometry core.

## Author contributions

H-CC, NM, KS, ER, CK, FB, LOB, ARF, and AG executed experiments. CMS analyzed TCGA datasets, FJ performed image analyses, and SW and NA produced Antibody Drug conjugates. DS provided linker technology. OG, IZ, SJA, MH, RM, ES, JDB, AG-G, SER and BH provided human healthy and tumour tissues, tumour microarrays, and patient-derived xenografts. Monoclonal antibodies against human LGR5 were generated by KS, MS, MarcdlR, and MaikedlR. MaikedlR and MarcdlR conceived the project, designed the experiments, analyzed results and wrote the manuscript.

## Data and materials availability

All data are available in the main text or the supplementary materials.

## References and Notes

1. Van de Wetering M, Sancho E, Verweij C, De Lau W, Oving I, Hurlstone A, Van der Horn K, Batlle E, Coudreuse D, Haramis AP, Tjon-Pon-Fong M, Moerer P, Van den Born M, Soete G, Pals S, Eilers M, Medema R, Clevers H. The β-catenin/TCF-4 complex imposes a crypt progenitor phenotype on colorectal cancer cells. Cell. 2002;111(2):241–50.

2. Koo BK, Clevers H. Stem cells marked by the r-spondin receptor LGR5. Gastroenterology. 2014;147(2):289–302.

3. Barker N, Huch M, Kujala P, van de Wetering M, Snippert HJ, van Es JH, Sato T, Stange DE, Begthel H, van den Born M, Danenberg E, van den Brink S, Korving J, Abo A, Peters PJ, Wright N, Poulsom R, Clevers H. Lgr5+ve Stem Cells Drive Self-Renewal in the Stomach and Build Long-Lived Gastric Units In Vitro. Cell Stem Cell. 2010;6(1):25–36.

4. Hugo J Snippert, Andrea Haegebarth, Maria Kasper, Viljar Jaks, Johan H van Es, Nick Barker, Marc van de Wetering, Maaike van den Born, Harry Begthel, Robert G Vries, Daniel E Stange, Rune Toftgård HC. Lgr6 Marks Stem Cells in the Hair Follicle That Generate All Cell Lineages of the Skin. Science. 2010;327(March):1385–9.

5. Trejo CL, Luna G, Dravis C, Spike BT, Wahl GM. Lgr5 is a marker for fetal mammary stem cells, but is not essential for stem cell activity or tumorigenesis. npj Breast Cancer. 2017;3(1):1–10.

6. Barker N, Rookmaaker MB, Kujala P, Ng A, Leushacke M, Snippert H, van de Wetering M, Tan S, Van Es JH, Huch M, Poulsom R, Verhaar MC, Peters PJ, Clevers H. Lgr5+ve Stem/Progenitor Cells Contribute to Nephron Formation during Kidney Development. Cell Rep. 2012;2(3):540–52.

7. Huch M, Dorrell C, Boj SF, Van Es JH, Li VSW, Van De Wetering M, Sato T, Hamer K, Sasaki N, Finegold MJ, Haft A, Vries RG, Grompe M, Clevers H. In vitro expansion of single Lgr5 + liver stem cells induced by Wnt-driven regeneration. Nature. 2013;494(7436):247–50.

8. Yamamoto Y, Sakamoto M, Fujii G, Tsuiji H, Kanetaka K, Asaka M, Hirohashi S. Overexpression of orphan G-protein-coupled receptor, Gpr49, in human hepatocellular carcinomas with β-catenin mutations. Hepatology. 2003;37(3):528–33.

9. McClanahan T, Koseoglu S, Smith K, Grein J, Gustafson E, Black S, Kirschmeier P, Samatar AA. Identification of overexpression of orphan G Protein-Coupled Receptor GPR49 in human colon and ovarian primary tumors. Cancer Biol Ther. 2006;5(4):419–26.

10. Tanese K, Fukuma M, Yamada T, Mori T, Yoshikawa T, Watanabe W, Ishiko A, Amagai M, Nishikawa T, Sakamoto M. G-protein-coupled receptor GPR49 is up-regulated in basal cell carcinoma and promotes cell proliferation and tumor formation. Am J Pathol. 2008;173(3):835–43.

11. Hagerling C, Owyong M, Sitarama V, Wang CY, Wang CY, Lin C, Van Den Bijgaart RJE, Van Den Bijgaart RJE, Koopman CD, Koopman CD, Koopman CD, Brenot A, Brenot A, Nanjaraj A, Wärnberg F, Jirström K, Klein OD, Klein OD, Werb Z, Plaks V, Plaks V. LGR5 in breast cancer and ductal carcinoma in situ: A diagnostic and prognostic biomarker and a therapeutic target. BMC Cancer. 2020;20(1):1–14.

12. Nakata S, Campos B, Bageritz J, Lorenzo Bermejo J, Becker N, Engel F, Acker T, Momma S, Herold-Mende C, Lichter P, Radlwimmer B, Goidts V. LGR5 is a marker of poor prognosis in glioblastoma and is required for survival of brain cancer stem-like cells. Brain Pathol. 2013;23(1):60–72.

13. Cosgun KNN, Hecht A, Yang X, Mangolini M, Aghajanirefah A, Chen Z, Xiao G, Klemm L, Hong C, Geng H, Jumaa H, Clevers H, Muschen M. Lgr5 Functions As Negative Regulator of Wnt Signaling in B Cells and Is Critical for Self-Renewal of Normal and Transformed B Cells. Blood. 2017;130(Supplement 1):3989.

14. Cosgun KN, Robinson M, Deb G, Yang X, Xiao G, Sadras T, Lee J, Chan L, Kume K, Mangolini M, Winchester J, Chen Z, Yang L, Geng H, Izraeli S, Song J, Chan WC, Polson A, Jumaa H, Clevers H, Müschen M. Lgr5-mediated restraint of β-catenin is essential for B-lymphopoiesis and leukemia-initiation. Prepr - Bioarchive. 2020;

15. Morgan RG, Mortensson E, Williams AC. Targeting LGR5 in Colorectal Cancer: Therapeutic gold or too plastic? Br J Cancer. 2018;118(11):1410–8.

16. Shimokawa M, Ohta Y, Nishikori S, Matano M, Takano A, Fujii M, Date S, Sugimoto S, Kanai T, Sato T. Visualization and targeting of LGR5 + human colon cancer stem cells. Nature. 2017;545(7653):187–92.

17. Kobayashi S, Yamada-Okabe H, Suzuki M, Natori O, Kato A, Matsubara K, Chen YJ, Yamazaki M, Funahashi S, Yoshida K, Hashimoto E, Watanabe Y, Mutoh H, Ashihara M, Kato C, Watanabe T, Yoshikubo T, Tamaoki N, Ochiya T, Kuroda M, Levine AJ, Yamazaki T. LGR5-positive colon cancer stem cells interconvert with drug-resistant LGR5-negative cells and are capable of tumor reconstitution. Stem Cells. 2012;30(12):2631–44.

18. Drago JZ, Modi S, Chandarlapaty S. Unlocking the potential of antibody–drug conjugates for cancer therapy. Nat Rev Clin Oncol. 2021;18(6):327–44.

19. Waldman AD, Fritz JM, Lenardo MJ. A guide to cancer immunotherapy: from T cell basic science to clinical practice. Nat Rev Immunol. 2020;20(11):651–68.

20. Slaney CY, Wang P, Darcy PK, Kershaw MH. CARs versus biTEs: A comparison between T cell– redirection strategies for cancer treatment. Cancer Discov. 2018;8(8):924–34.

21. Snyder JC, Rochelle LK, Ray C, Pack TF, Bock CB, Lubkov V, Lyerly HK, Waggoner AS, Barak LS, Caron MG. Inhibiting clathrin-mediated endocytosis of the leucine-rich G protein-coupled receptor-5 diminishes cell fitness. J Biol Chem. 2017;292(17):7208–22.

22. Rossmann M, Greive SJ, Moschetti T, Dinan M, Hyvönen M. Development of a multipurpose scaffold for the display of peptide loops. Protein Eng Des Sel. 2017;30(6):419–30.

23. Chen P han, Chen X, Lin Z, Dev G, Chen P han, Chen X, Lin Z, Fang D, He X. The structural basis of R-spondin recognition by LGR5 and RNF43 Email alerting service The structural basis of R-spondin recognition by LGR5 and RNF43. 2013;1345–50.

24. Korinek V, Barker N, Morin PJ, Van Wichen D, De Weger R, Kinzler KW, Vogelstein B, Clevers H. Constitutive transcriptional activation by a β-catenin-Tcf complex in APC(-/-) colon carcinoma. Science. 1997;275(5307):1784–7.

25. Junttila MR, Mao W, Wang X, Wang BE, Pham T, Flygare J, Yu SF, Yee S, Goldenberg D, Fields C, Eastham-Anderson J, Singh M, Vij R, Hongo JA, Firestein R, Schutten M, Flagella K, Polakis P, Polson AG. Targeting LGR5+ cells with an antibody-drug conjugate for the treatment of colon cancer. Sci Transl Med. 2015;7(314):1–12.

26. Gong X, Azhdarinia A, Ghosh SC, Xiong W, An Z, Liu Q, Carmon KS. LGR5-Targeted antibody-drug conjugate eradicates gastrointestinal tumors and prevents recurrence. Mol Cancer Ther. 2016;15(7):1580–90.

27. Nguyen HG, Blank A, Dawson HE, Lugli A, Zlobec I. Classification of colorectal tissue images from high throughput tissue microarrays by ensemble deep learning methods. Sci Rep. 2021;11(1):1–11.

28. Zucman-rossi J, Villanueva A, Nault J charles, Llovet JM. Genetic Landscape and Biomarkers of Hepatocellular Carcinoma. YGAST. 2015;149(5):1226–1239.e4.

29. Snyder JC, Rochelle LK, Lyerly HK, Caron MG, Barak LS. Constitutive internalization of the Leucine-rich g protein-coupled receptor-5 (LGR5) to the trans-Golgi network. J Biol Chem. 2013;288(15):10286–7.

30. Carmon KS, Gong X, Yi J, Wu L, Thomas A, Moore CM, Masuho I, Timson DJ, Martemyanov KA, Liu QJ. LGR5 receptor promotes cell-cell adhesion in stem cells and colon cancer cells via the IQGAP1-Rac1 pathway. J Biol Chem. 2017;292(36):14989–5001.

31. Kurten RC, Cadena DL, Gill GN. Enhanced degradation of EGF receptors by a sorting nexin, SNX1. Science (80-). 1996;272(5264):1008–10.

32. van Dam EM, Stoorvogel W. Dynamin-dependent Transferrin Receptor Recycling by Endosome-derived Clathrin-coated Vesicles. Mol Biol Cell. 2002;13(January):169–182.

33. Bargh JD, Walsh SJ, Isidro-Llobet A, Omarjee S, Carroll JS, Spring DR. Sulfatase-cleavable linkers for antibody-drug conjugates. Chem Sci. 2020;11(9):2375–80.

34. Walsh SJ, Omarjee S, Galloway WRJD, Kwan TTL, Sore HF, Parker JS, Hyvönen M, Carroll JS, Spring DR. A general approach for the site-selective modification of native proteins, enabling the generation of stable and functional antibody-drug conjugates. Chem Sci. 2019;10(3):694–700.

35. Zhong XS, Matsushita M, Plotkin J, Riviere I, Sadelain M. Chimeric antigen receptors combining 4-1BB and CD28 signaling domains augment PI 3 kinase/AKT/Bcl-X L activation and CD8 T cell-mediated tumor eradication. Mol Ther. 2010;18(2):413–20.

36. Siegler EL, Zhu Y, Wang P, Yang L. Off-the-Shelf CAR-NK Cells for Cancer Immunotherapy. Cell Stem Cell. 2018;23(2):160–1.

37. Mitwasi N, Feldmann A, Arndt C, Koristka S, Berndt N, Jureczek J, Loureiro LR, Bergmann R, Máthé D, Hegedüs N, Kovács T, Zhang C, Oberoi P, Jäger E, Seliger B, Rössig C, Temme A, Eitler J, Tonn T, Schmitz M, Hassel JC, Jäger D, Wels WS, Bachmann M. “UniCAR”-modified off-the-shelf NK-92 cells for targeting of GD2-expressing tumour cells. Sci Rep. 2020;10(1):1–16.

38. Leung C, Tan SH, Barker N. Recent Advances in Lgr5+ Stem Cell Research. Trends Cell Biol. 2018;28(5):380–91.

39. Haegebarth A, Clevers H. Wnt signaling, Lgr5, and stem cells in the intestine and skin. Am J Pathol. 2009;174(3):715–21.

40. Nusse R, Clevers H. Wnt/β-Catenin Signaling, Disease, and Emerging Therapeutic Modalities. Cell. 2017;169(6):985–99.

41. Berraondo P, Ochoa MC, Olivera I, Melero I. Immune desertic landscapes in hepatocellular carcinoma shaped by β-catenin activation. Cancer Discov. 2019;9(8):1003–5.

42. Galarreta MR De, Bresnahan E, Molina-sánchez P, Lindblad KE, Maier B, Sia D, Puigvehi M, Miguela V, Casanova-acebes M, Dhainaut M, Villacorta-martin C, Singhi AD, Moghe A, Felden J Von, Grinspan LT, Wang S, Kamphorst AO, Monga SP, Brown BD, Villanueva A, Llovet JM, Merad M, Lujambio A. β-catenin activation promotes immune escape and resistance to anti-PD-1 therapy in hepatocellular carcinoma. 2019;9:1124–41.

43. Yau T, Park J won, Finn RS, Cheng A lii, Mathurin P, Edeline J, Kudo M, Harding JJ, Merle P, Abou-alfa GK, El-khoueiry AB, Melero I, Begic D, Chen G, Neely J, Wisniewski T, Tschaika M, Sangro B. Nivolumab versus sorafenib in advanced hepatocellular carcinoma (CheckMate 459): a randomised, multicentre, open-label, phase 3 trial. Lancet Oncol. 2022;23(1):77–90.

44. De La Roche M, Ibrahim AEK, Mieszczanek J, Bienz M. LEF1 and B9L shield b-catenin from inactivation by axin, desensitizing colorectal cancer cells to tankyrase inhibitors. Cancer Res. 2014;74(5):1495–505.

45. Chen H, Eling N, Martinez-Jimenez CP, O’Brien LM, Carbonaro V, Marioni JC, Odom DT, Roche M. IL-7-dependent compositional changes within the γδ T cell pool in lymph nodes during ageing lead to an unbalanced anti-tumour response. EMBO Rep. 2019;20(8):1–17.

46. James JR. Tuning ITAM multiplicity on T cell receptors can control potency and selectivity to ligand density. Sci Signal. 2018;11(531).

47. Roovers RC, Herpers B, James M, Eppink B, Cortina C, Maussang-Detaille D, Kolfschoten I, Boy SF, van de Wetering M, de Lau W, Doornbos R, Clements C, Basmeleh A, Bartelink W, van de Zande VZ, Yan K, Salinaro L, Bakker L, de Kruif J, Clevers H, Vries R, Batlle E, Price L, Throsby M. Identification and characterisation of MCLA-158 as an anti-EGFRxLGR5 bispecific antibody that potently inhibits patient-derived CRC organoid growth. Am Assoc Cancer Res. 2017;158.

48. Inglis DJ, Licari J, Georgiou, Kristen R. Wittwer NL, Hamilton RW, Beaumont, Donna M. Scherer MA, Lavranos TC. Abstract 3910: Characterization of BNC101 a human specific monoclonal antibody targeting the GPCR LGR5: First-in-human evidence of target engagement. In: American Association for Cancer Research, editor. Proceedings of the American Association for Cancer Research Annual Meeting 2018. Cancer Research; 2018.

