## Supplemental Material for "LGR5 targeting molecules as therapeutic agents for multiple cancer types"

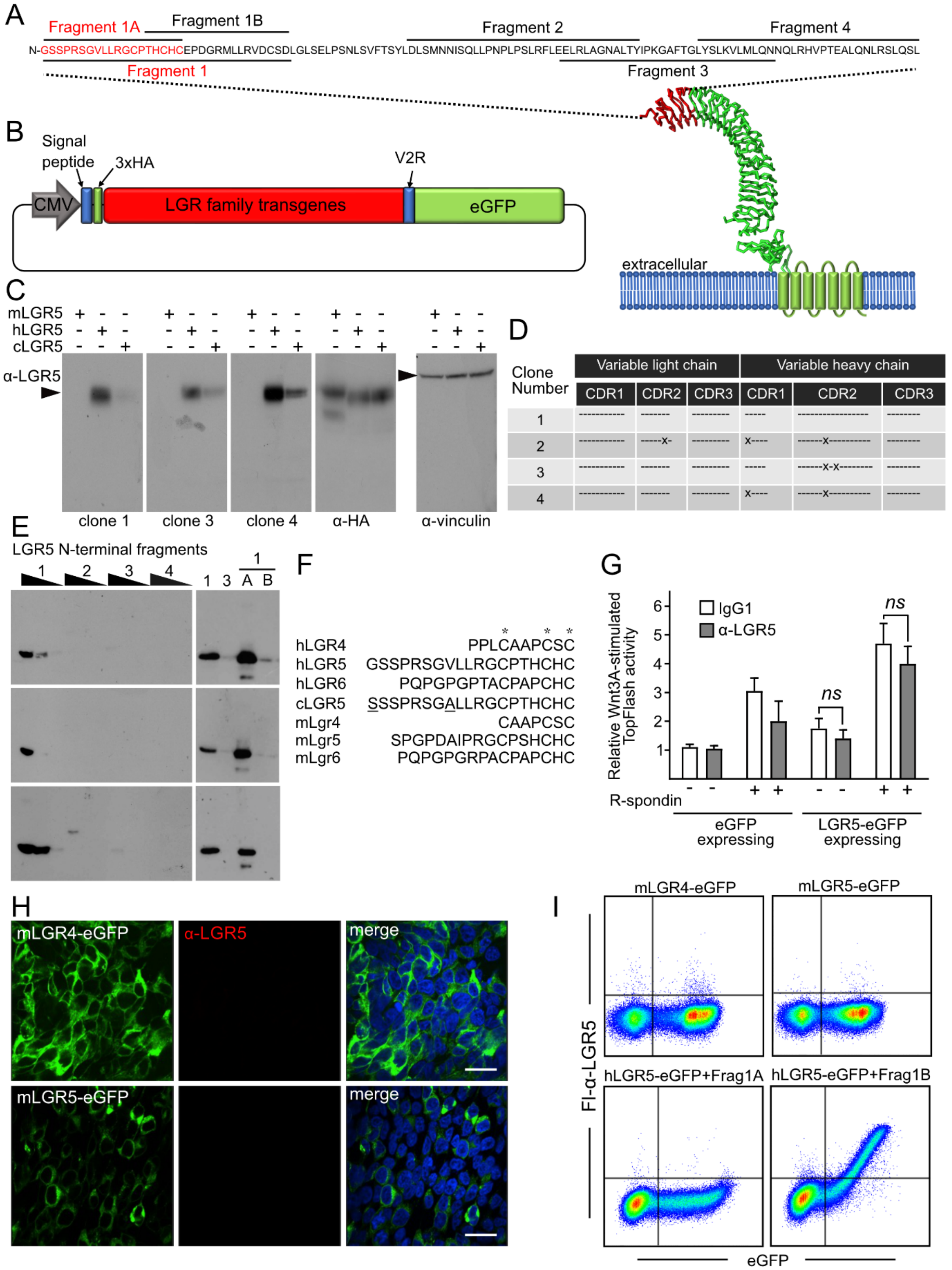


| Clone Number | VL CDR1 | VL CDR2 | VL CDR3 | VH CDR1 | VH CDR2 | VH CDR3 |
|---|---|---|---|---|---|---|
| 1 | ---------- | ------- | --------- | ----- | ---------------- | -------- |
| 2 | ---------- | -----x- | --------- | x---- | ------x--------- | -------- |
| 3 | ---------- | ------- | --------- | ----- | ------x-x------- | -------- |
| 4 | ---------- | ------- | --------- | x---- | ------x--------- | -------- |

**Supplemental Figure 1. Specificity of LGR5 antibodies generated in the study.**

A. Amino acid sequence of the human LGR5 antigen used for mouse immunization and generation of $\alpha$-LGR5; numbering starts at Gly1 in the processed hLGR5, lacking the signal sequence. The sequence is annotated with the Fragments used in the RAD display experiments that map the $\alpha$-LGR5 epitope to Frag1A. *Below* – location of the antigenic region (in red) within the structure of the extracellular domain of LGR5 (atomic coordinates for the model taken from (1).
B. Configuration of the LGR family transgenic constructs used in the study. All expressed LGR proteins contain a common N-terminal hemagglutinin (HA) tag and fusion at the C-terminus to the vasopressin V2 receptor C-terminal tail (V2R) followed by eGFP.
C. Western blot analysis of HEK293T lysates expressing the human LGR family transgenes probed with $\alpha$-LGR5 hybridoma clones 1, 3 and 4 and antibodies to HA and vinculin, as noted. No specific immune reactivity was observed when probing the western blots with the other 14 hybridoma clones.
D. Sequence conservation amongst the $\alpha$-LGR5 hybridoma clones within the complementary determining regions (CDRs). Conserved amino acids relative to $\alpha$-LGR5 clone 1 for clones 2-4 are represented by a dash. Amino acid differences are shown with a closed circle.
E. Western blot analysis of the Fragments delineated above (Suppl. Fig. 1A) as RAD-displayed fusion peptides using $\alpha$-LGR5 hybridoma clones 1, 3 and 4.
F. Sequence alignment of the N-terminal 15 amino acids of human LGR5, corresponding to Frag1A, with the corresponding region in the other LGR family members. Sequences were aligned based on three invariant cysteine residues denoted by asterisks. The amino acid difference in the *cynomolgus* sequence is underlined.
G. Wnt pathway reporter assays (TopFlash assays) for HEK293T cells transfected with either eGFP or human LGR5-eGFP (hLGR5-eGFP), treated with Wnt3A ligand, R-spondin and either IgG1 or $\alpha$-LGR5 at levels of approximately 10-fold molar excess over Wnt3A ligand. ns, no significant difference, determined by two-tailed t-test. Error bars indicate standard deviation (SD) for 3 biological replicates. *ns*, no significant difference in Wnt pathway reporter activity.
H. Immunofluorescent detection of HEK293T cells expressing transgenic LGR4-eGFP or LGR5-eGFP (*left panels*, green) using Fl-$\alpha$-LGR5 (*middle panels*, red). *Right panels* merged fluorescent signals. Scale bars, 10 $\mu$M.
I. Flow cytometric analysis of HEK293T cells expressing mLGR4-eGFP (*top left*), mLGR5-eGFP (*top right*) and hLGR5-eGFP (*bottom panels*) using Fl-$\alpha$-LGR5. For analysis of the hLGR5-eGFP expressing HEK293T cells, Fl-$\alpha$-LGR5 was pre-incubated with either RAD-Frag1A or RAD-Frag1B (*bottom left and right*).

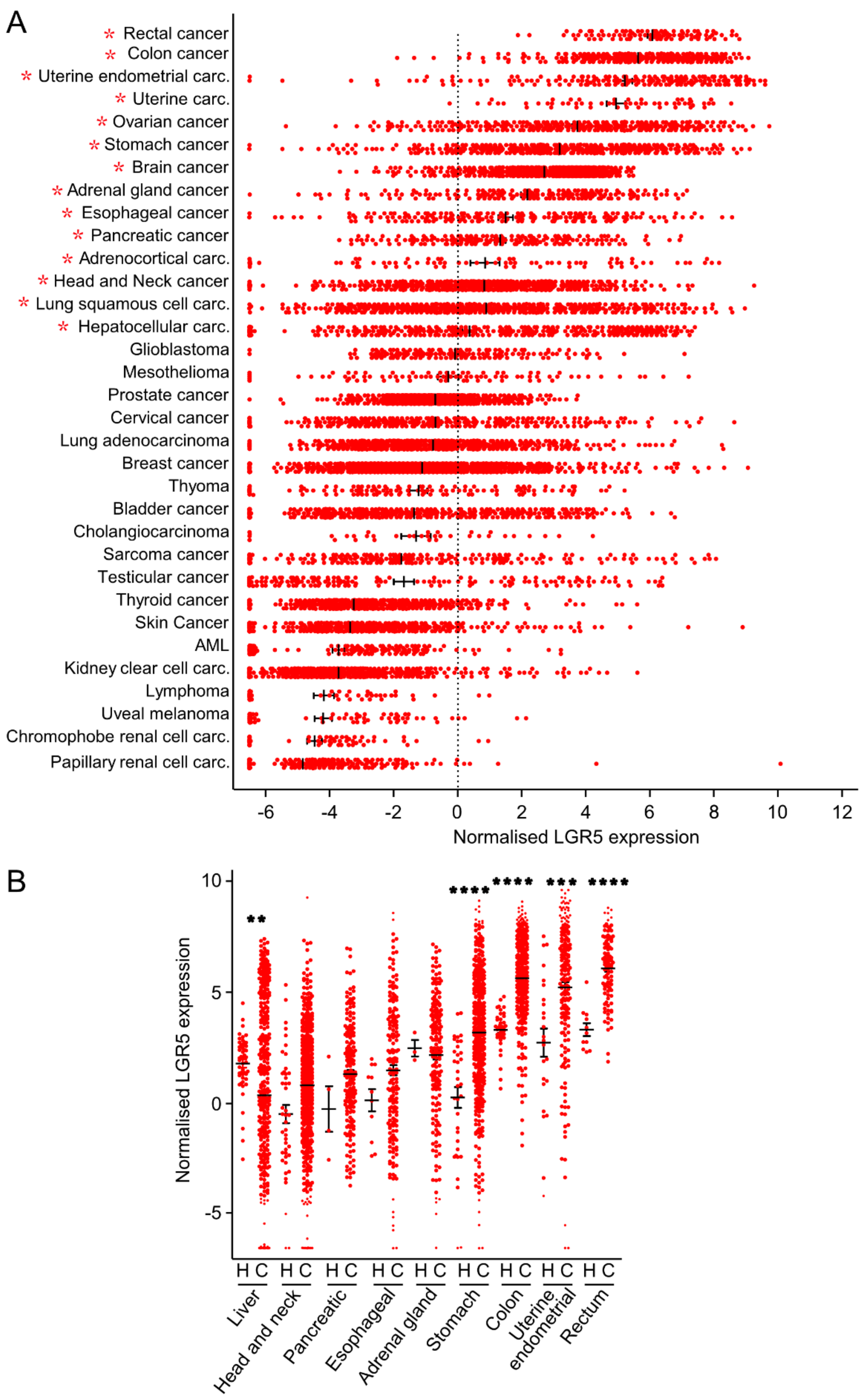


Supplemental Figure 2 (page1)

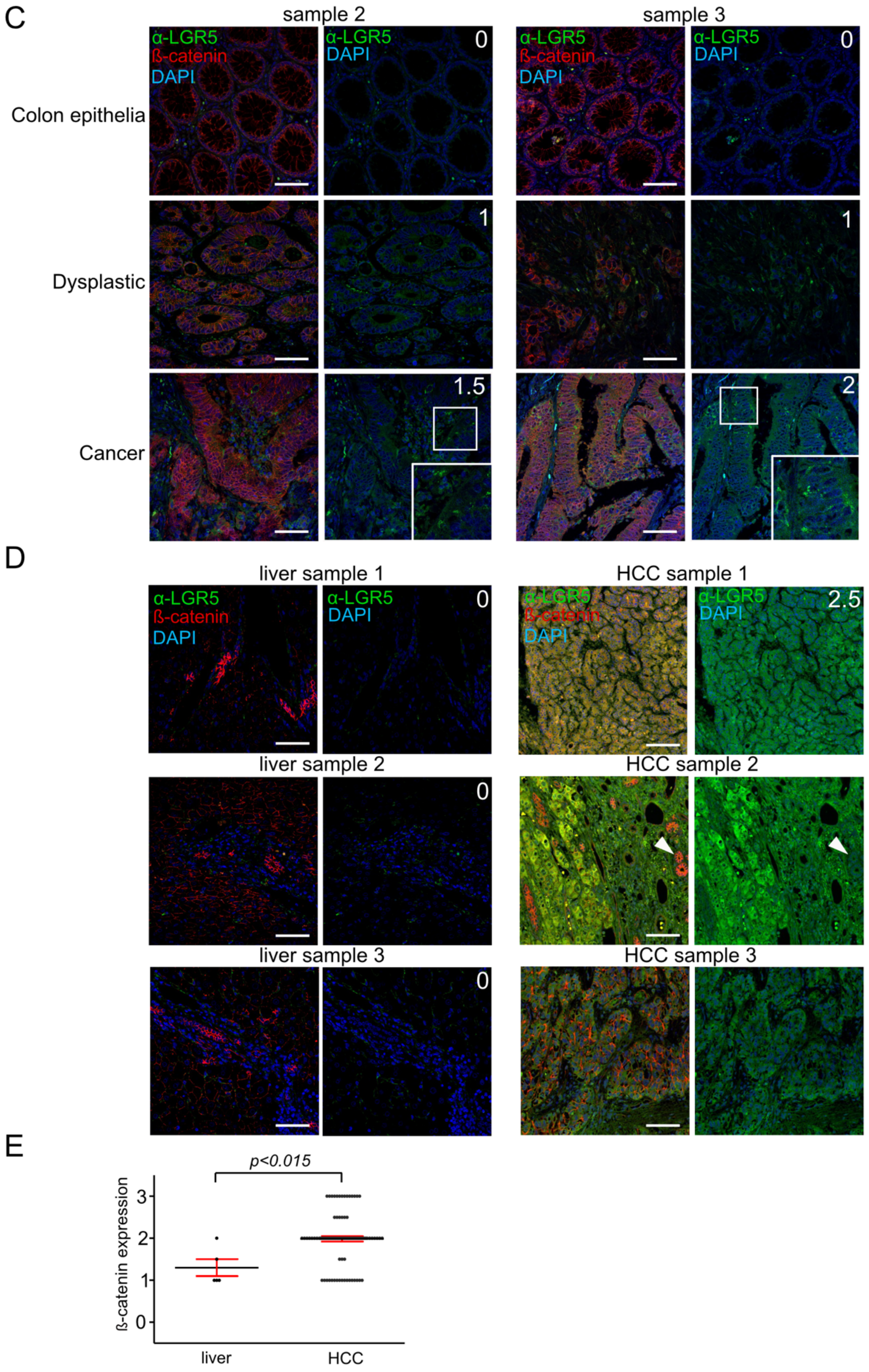


Supplemental Figure 2 (page 2)

F

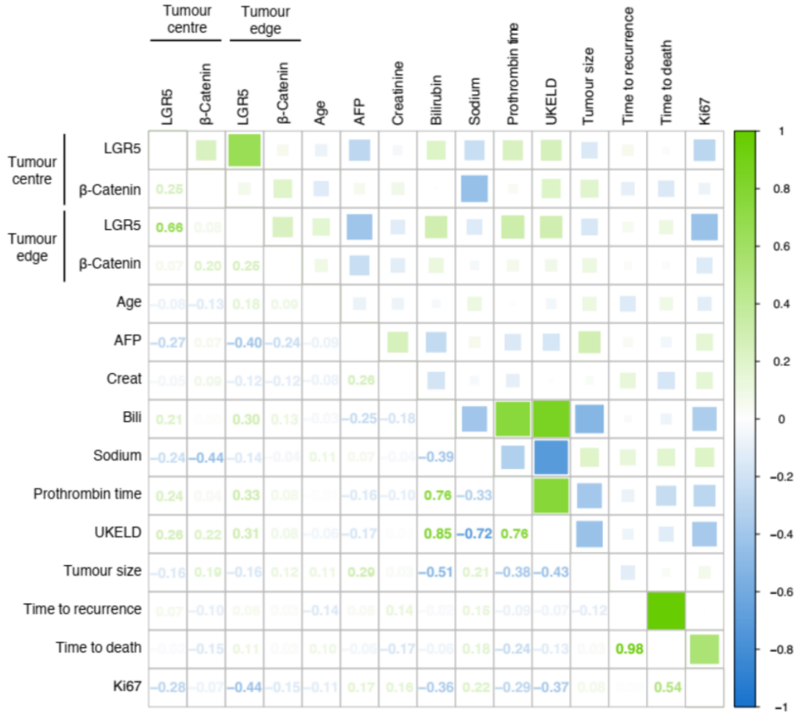


G

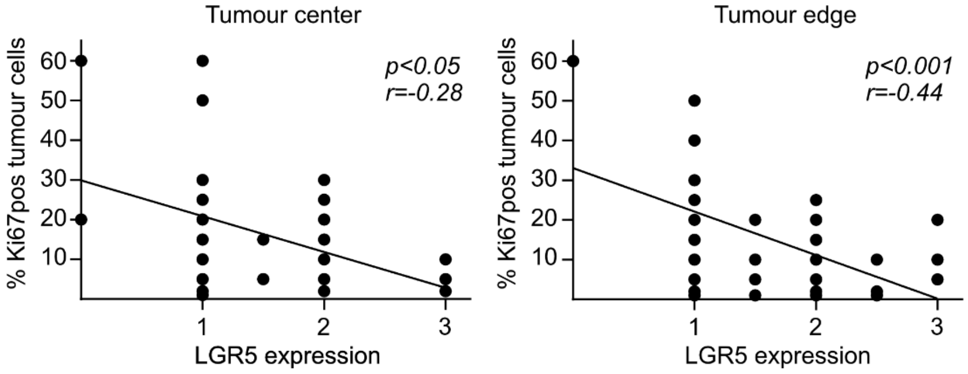


Supplemental Figure 2 (page 3)

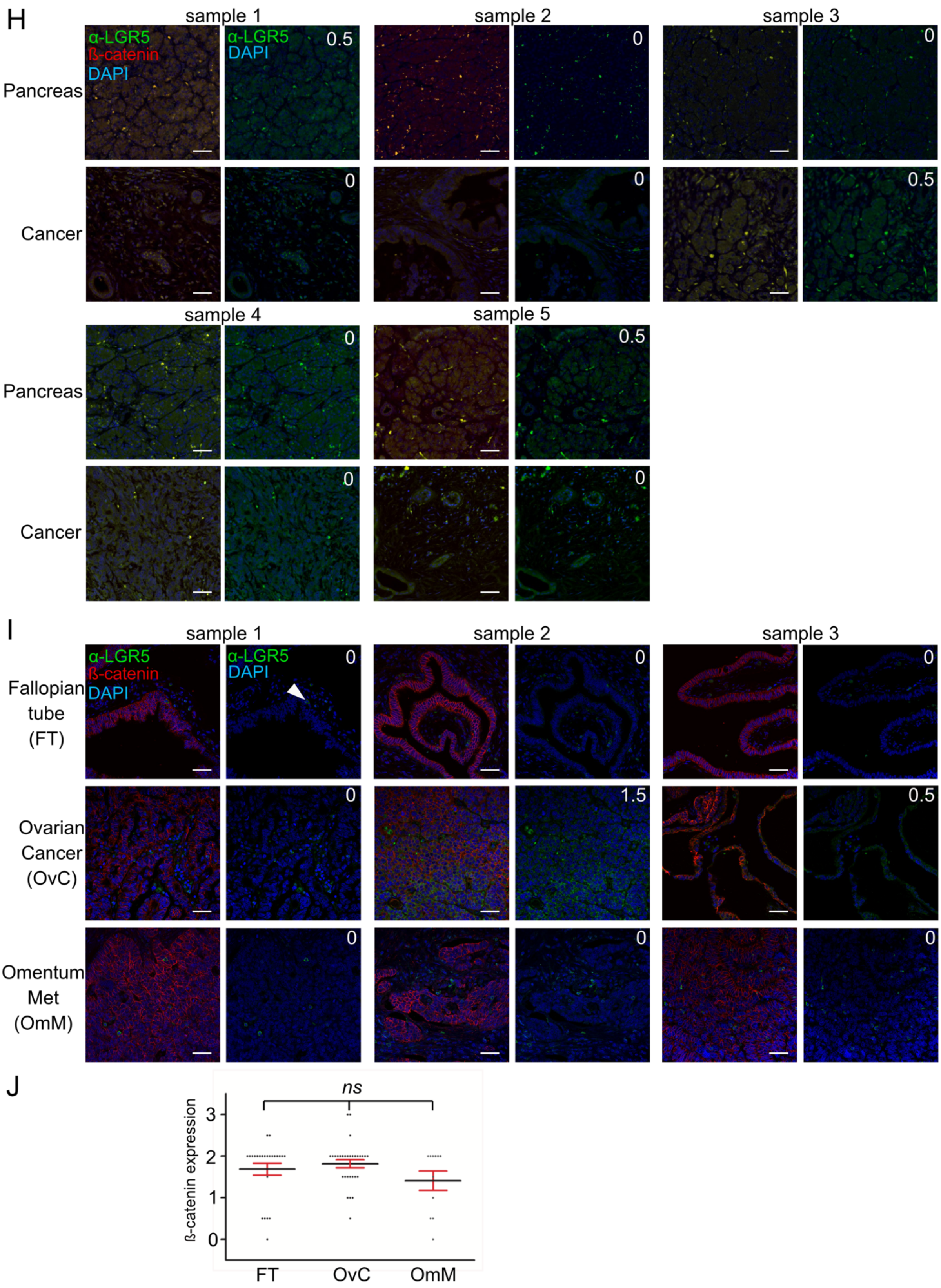


Supplemental Figure 2 (page 4)

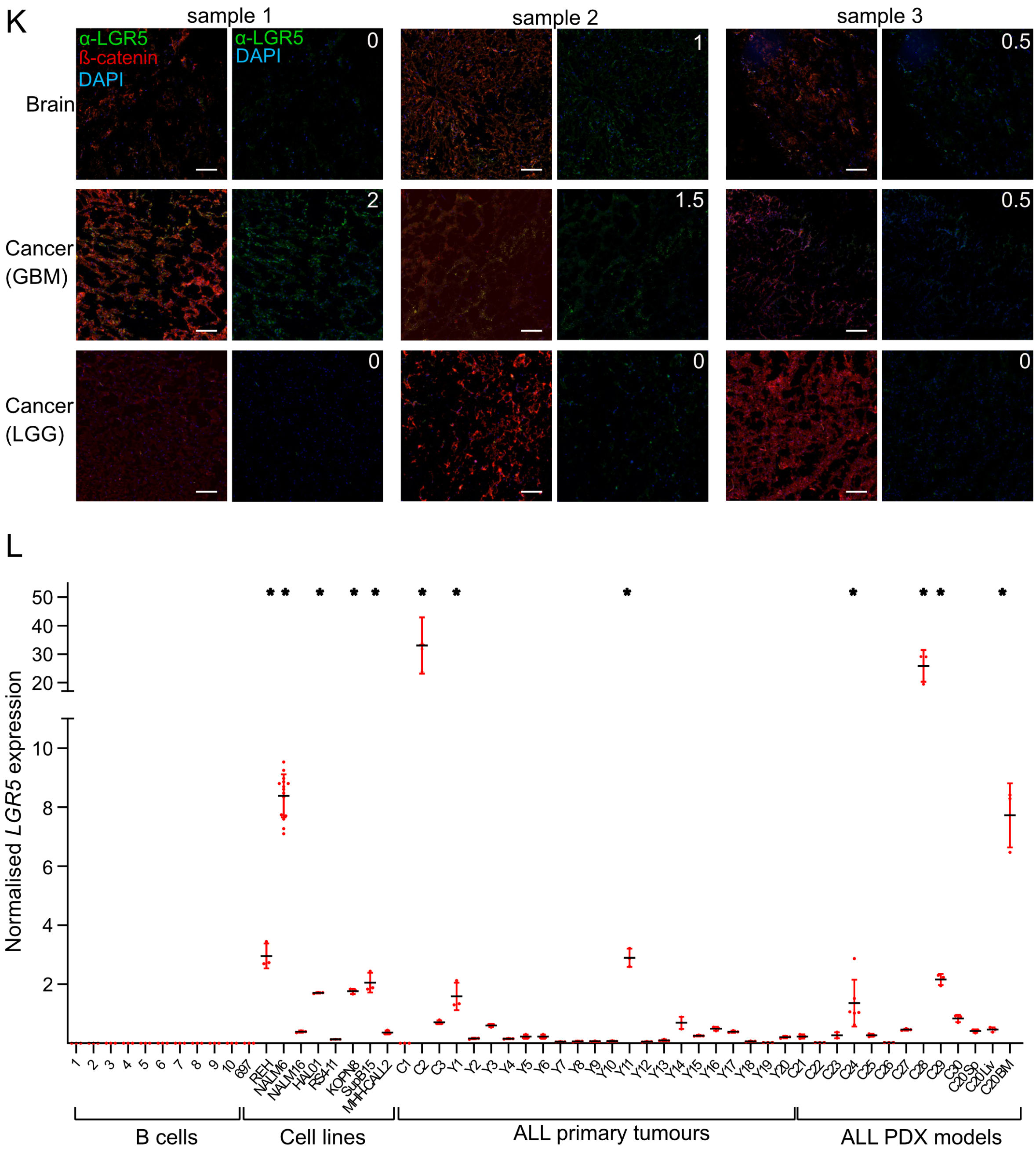


Supplemental Figure 2 (page 5)

**Supplemental Figure 2. LGR5 expression analysis in cancers**

A. Normalised (log2 median-centred) LGR5 gene expression levels for cancer subtypes, ordered by median LGR5 gene expression. Read counts were quantile normalized across the genome for direct comparison amongst cancer subtypes and sample sets and median expression levels for extracted LGR5 data determined across the entire dataset. The dotted line indicates the median LGR5 expression across all pan-cancer tumours. Tumour subtypes for which more than 70% of samples had higher than median LGR5 expression (dashed line) were defined as "high LGR5 tumours" are denoted with a red *.

B. Comparison of LGR5 gene expression between healthy tissue (H) and cancer (C) for selected high LGR5 tumours. Significance difference in LGR5 expression between cancer and healthy tissue (Wilcoxon test) are indicated: **, *p<0.01*, ***, *p<0.001*, **** *p<0.0001*.

C. LGR5 and β-catenin protein levels in sections from two CRC tumour resections with regions of normal colon epithelia (*top panels*), dysplastic tissue (*middle panels*) and cancer (*lower panels*). White numbers are the relative values for LGR5 protein expression using the scoring system applied to all tissue and cancer biopsies. Scale bars, 10 μM.

D. Representative images from 3 liver samples (*left panels*) and 3 of the 95 HCC cases from the Cambridge HCC TMA (*right panels*). White numbers are the score for LGR5 expression levels. Scale bars, 10 μM.

E. Quantitation of β-catenin expression levels in 8 liver resections (liver) and the HCC cases from the Cambridge HCC TMA. Level of significant difference, *p-value*, between the sample sets was determined by two-tailed t-test.

F. Correlation matrix between LGR5 or β-catenin protein expression levels and phenotypic metrics determined for the biopsies comprising the Cambridge HCC TMA. AFP, levels of the tumour serum biomarker alpha-fetoprotein at the time of transplant; UKELD, UK Model for End-Stage Liver Disease score (2), Ki67, quantification of Ki67 immunohistochemistry antigen used as an index of proliferation. Correlation values were computed pairwise using complete observations (i.e., removing missing values) using the Spearman Rank method (values shown left of the diagonal). Level of correlation amongst phenotypic metrics is scaled from highly positive - dark green colour and large squares, to highly negative - dark blue and large squares (shown to the right of the diagonal. Plot of Ki67 levels versus LGR5 protein expression in the tumour centre (left graph) or the tumour edge (right edge) for HCC cases with clinical and molecular features of the non-proliferative HCC sub-class. Significance for inverse correlation of LGR5 expression levels and percent cells expressing Ki67 determined by Spearman correlation test and represented by p-value (*p*) and correlation coefficient (*r*).

G. Images of β-catenin and LGR5 expression in the five matched pancreas samples and pancreatic cancer cases (Cancer). White numbers, scored values for relative LGR5 protein expression. Low level expression of the two proteins was apparent for all samples. Scale bars, 40 μM.

H. Representative images for fallopian tube tissue (FT; *left panel set*), ovarian cancers (OvC; *middle panel set*) and omentum cancers (OmC; *right panel set*). Arrowhead shown on the first fallopian tube sample indicate a very rare instance of epithelial cells containing LGR5 positive intracellular puncta. White numbers, scored values for relative LGR5 protein expression. Scale bars, 40 μM.

I. Relative expression levels of β-catenin in the fallopian tube, ovarian cancer and omentum cancer sample sets. There was no significance in β-catenin protein levels amongst fallopian tube (FT), ovarian cancer (OvC) or omentum cancer (OmC) samples, determined using two-tailed t-test.

J. Representative images for brain tissue (*left panel set*), GBM (*middle panel set*) and LGG (*right panel set*). White numbers, scored values for relative LGR5 expression. Scale bars, 40 μM.

K. Quantification of LGR5 transcript levels for samples (grouped in **Fig. 2H**) as follows – healthy donor B cells (B-cells; 10 samples), B-ALL cell lines (Cell lines; 9 lines), CD19-enriched cell populations from primary B-ALL cases (ALL primary; 15 samples) and CD19-enriched populations from B-ALL tumour cells maintained as PDX models (ALL-PDX 22 samples). Samples beginning with C refer to biopsies obtained from the Cambridge biobank, samples beginning with Y were from the York biobank (see Materials and Methods). Error bars represent SD for a minimum of three replicates per sample. Level of significance, *p-value*, was determined by two-tailed t-tests comparing LGR5 expression between individual samples and the average of all healthy donor B cells and shown at the *, *p<0.001* level of significance.

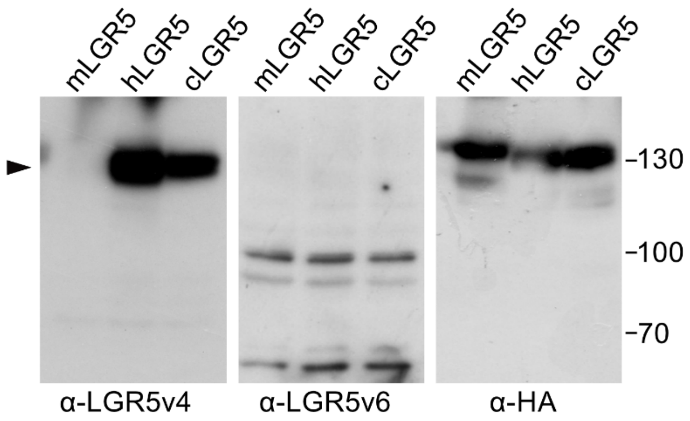


**Supplemental Figure 3. Validation of α-LGR5v4**

Western blot analysis of LGR5 protein levels in lysates from HEK293T cell overexpressing, *lanes 1-3*, mLGR5-eGFP, hLGR5-eGFP and cLGR5-eGFP using humanised α-LGR5v4, α-LGR5v6 and α-HA. Arrow denotes approximate migration distance of eGFP fusions.

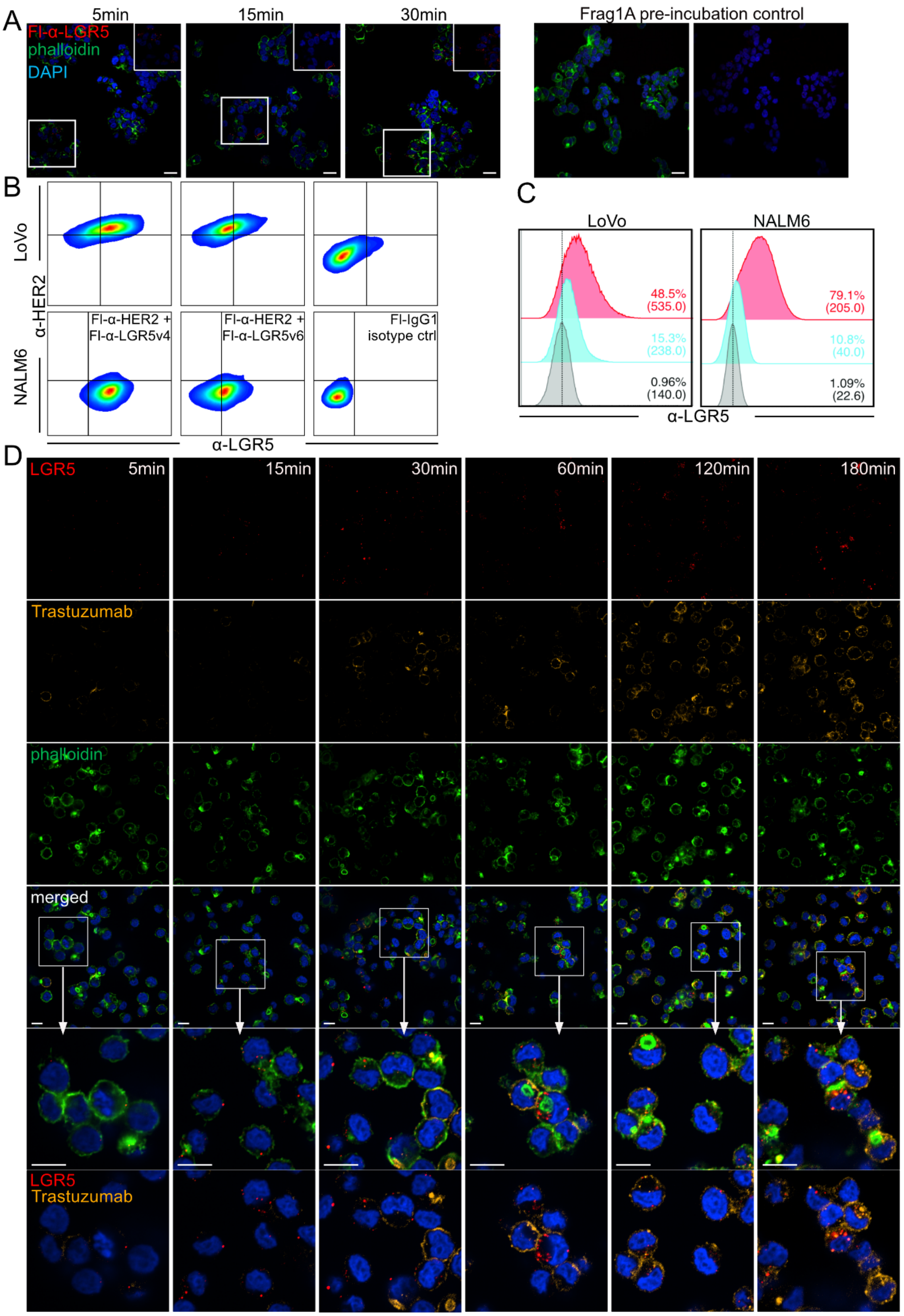
A
5min
15min
30min
Frag1A pre-incubation control
Fl-α-LGR5
phalloidin
DAPI
B
LoVo
NALM6
α-HER2
Fl-α-HER2 +
Fl-α-LGR5v4
Fl-α-HER2 +
Fl-α-LGR5v6
Fl-IgG1
isotype ctrl
α-LGR5
C
LoVo
NALM6
48.5%
(535.0)
15.3%
(238.0)
0.96%
(140.0)
79.1%
(205.0)
10.8%
(40.0)
1.09%
(22.6)
α-LGR5
D
LGR5
5min
15min
30min
60min
120min
180min
Trastuzumab
phalloidin
merged
LGR5
Trastuzumab

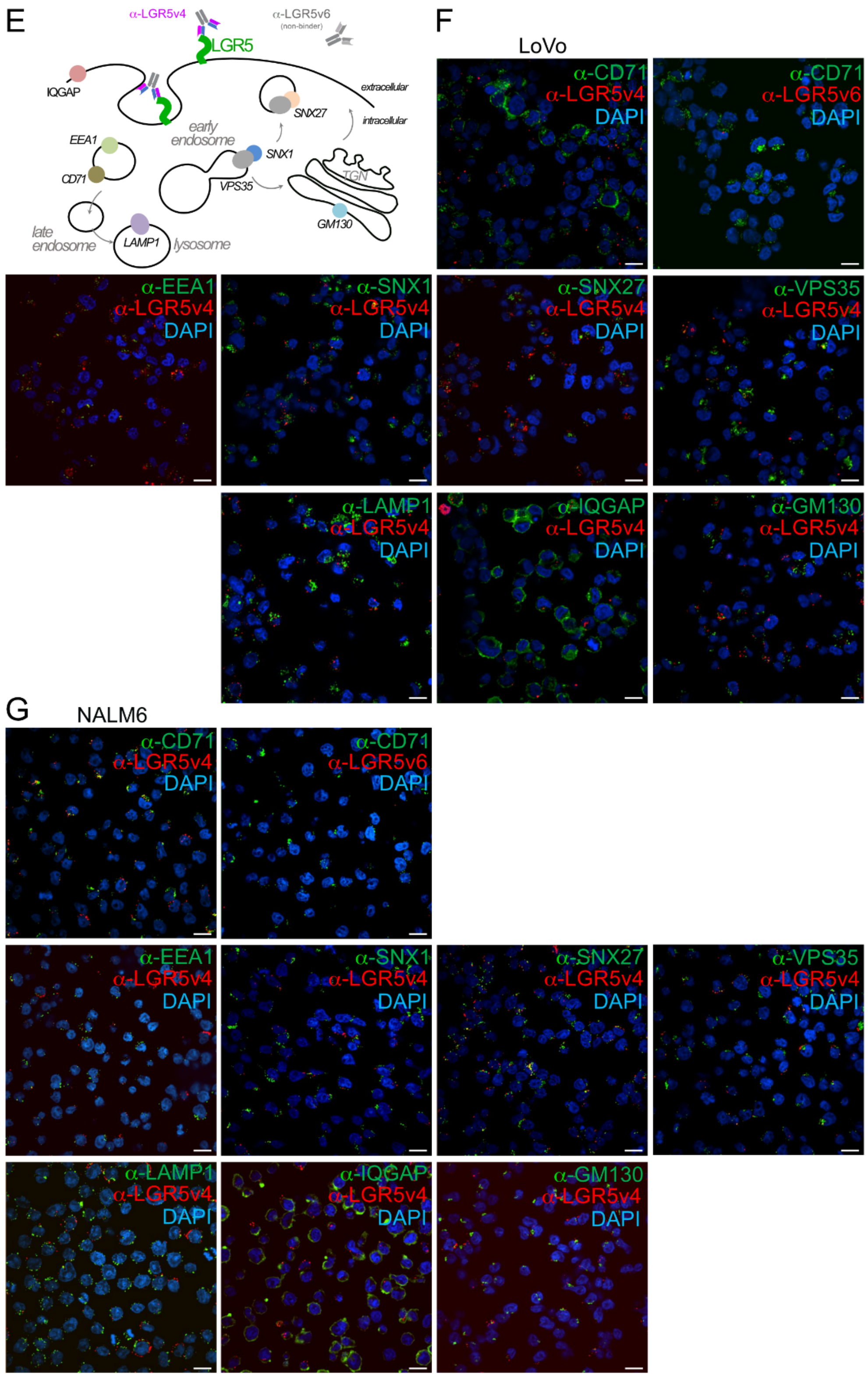


Supplemental Figure 4 (page 2)

**Supplemental Figure 4. LGR5 internalisation in NALM6 and LoVo cells.**

A. Time course of Fl-α-LGR5 (red) internalisation by LoVo cells. F-actin and nuclei were visualised by Alexa488-Phalloidin and Hoechst probes, respectively. For the two panels on the right, Fl-α-LGR5 was pre-incubated with Frag1A. In the rightmost image, the signal from Alexa488 phalloidin has been omitted. Scale bars, 10 μM.
B. Flow cytometric analysis of LoVo and NALM6 cells after 60 minutes co-incubation with Fl-α-LGR5v4 and Fl-α-HER2 (*left panels*), Fl-α-LGR5v4 and Fl-α-HER2 (middle panels) or Fl-IgG1 isotype controls.
C. Flow cytometric analyses of LoVo and NALM6 cells after 60 minutes incubation with Fl-α-LGR5v4 (red) or Fl-α-LGR5v4 pre-incubated with Frag1A (cyan) or Fl-IgG1 isotype control (grey).
D. Representative fields of view for time course of Fl-α-LGR5v4 and Fl-α-HER2 association and internalisation by NALM6 cells. Scored association and internalisation data is summarised in **Fig. 4D and E**. Scale bars, 20 μM.
E. Graphical representation of the intracellular markers used for immunofluorescence detection and co-localisation with internalised Fl-α-LGR5v4.
F. Representative single plane images showing immune detection of the intracellular markers (green) in LoVo cells that have been incubated with Fl-α-LGR5v4 (red) or Fl-α-LGR5v6 (second panel only) for 30 minutes. Scale bars, 10 μM.
G. Representative single plane images showing immune detection of the intracellular markers (green) in NALM6 cells that have been incubated with Fl-α-LGR5v4 (red) or Fl-α-LGR5v6 (second panel only) for 30 minutes. Scale bars, 10 μM.

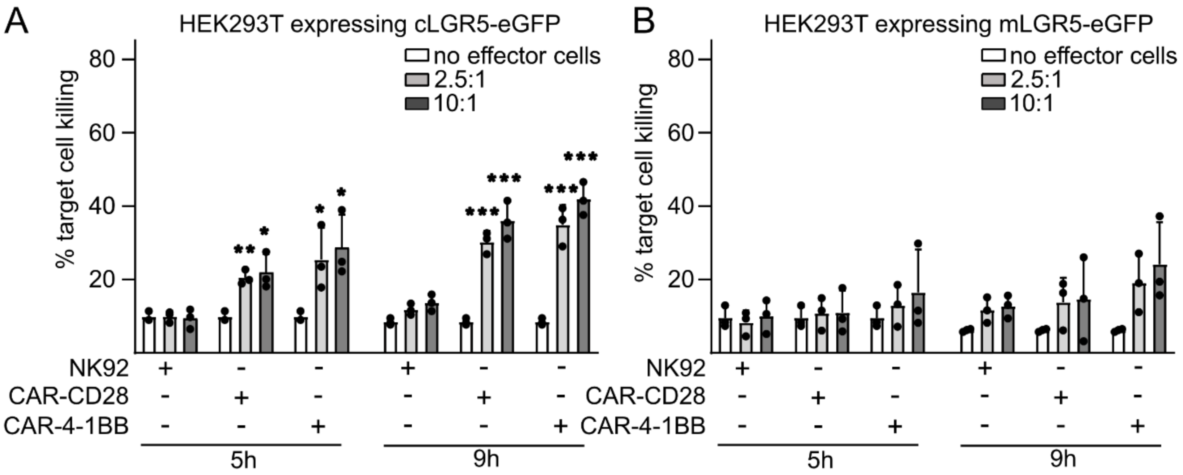


**Supplementary Figure 5. α-LGR5-mediated cell killing in BiTE and CAR modalities**

A. Cell killing activity of NK92 cells, or α-LGR5-CAR-CD28- or α-LGR5-CAR-4-1BB-expressing NK92 cells. Target cells were HEK293T cells overexpressing cLGR5-eGFP that were incubated with effector NK92 cells at effector to target ratios of 2.5:1 and 10:1 for 5h or 9h. Error bars represent SD for three independent experiments. Significant differences in target cell killing in the presence or absence of target cells was determined by two-tailed t-test and is shown at the *, *p<0.05* and **, *p<0.005* or **, *p<0.001* levels of significance. There were no significant differences in target cell killing with cellular ratios of 2.5:1 or 10:1.
B. Cell killing activity of NK92 cells, or α-LGR5-CAR-CD28- or α-LGR5-CAR-4-1BB-expressing NK92 cells. Target cells were HEK293T cells overexpressing mLGR5-eGFP that were incubated with effector NK92 cells at effector to target ratios of 2.5:1 and 10:1 for 5h or 9h. Error bars represent SD for three independent experiments. There were no significant differences in target cell killing activity amongst the experimental groups.

### Supplemental references


1. Peng WC, DeLau W, Forneris F, Granneman JCM, Huch M, Clevers H, et al. Structure of Stem Cell Growth Factor R-spondin 1 in Complex with the Ectodomain of Its Receptor LGR5. Cell Rep [Internet]. 2013;3(6):1885–92. Available from: http://dx.doi.org/10.1016/j.celrep.2013.06.009
2. Barber K, Madden S, Allen J, Collett D, Neuberger J, Gimson A. Elective liver transplant list mortality: Development of a United Kingdom end-stage liver disease score. Transplantation. 2011;92(4):469–76.
